# Nucleoporin 153 links nuclear pore complex to chromatin architecture by mediating CTCF and cohesin binding at *cis*-regulatory elements and TAD boundaries

**DOI:** 10.1101/2020.02.04.934398

**Authors:** Shinichi Kadota, Jianhong Ou, Yuming Shi, Jiayu Sun, Eda Yildirim

**Affiliations:** Department of Cell Biology, Duke Medical Center, Durham, NC 27710 USA; Duke Cancer Institute, Duke University, Durham, NC 27710 USA; Duke Regeneration Next Institute, Duke University, Durham, NC 27710 USA

**Keywords:** Nuclear pore complex, nucleoporin, chromatin structure, transcription, chromatin architecture, nuclear structure

## Abstract

The nuclear pore complex (NPC) components, nucleoporins (Nups), have been proposed to mediate spatial and temporal organization of chromatin during gene regulation. Nevertheless, we have little understanding on the molecular mechanisms that underlie Nup-mediated chromatin structure and transcription in mammals. Here, we show that Nucleoporin 153 (NUP153) interacts with the chromatin architectural proteins, CTCF and cohesin, and mediates their binding across *cis*-regulatory elements and TAD boundaries in mouse embryonic stem (ES) cells. NUP153 depletion results in altered CTCF and cohesin occupancy and differential gene expression. This function of NUP153 is most prevalent at the developmental genes that show bivalent chromatin state. To dissect the functional relevance of NUP153-mediated CTCF and cohesin binding during transcriptional activation or silencing, we utilized epidermal growth factor (EGF)-inducible immediate early genes (IEGs). We found that NUP153 binding at the *cis*-regulatory elements controls CTCF and cohesin binding and subsequent POL II pausing during the transcriptionally silent state. Furthermore, efficient and timely transcription initiation of IEGs relies on NUP153 and occurs around the nuclear periphery suggesting that NUP153 acts as an activator of IEG transcription. Collectively, these results uncover a key role for NUP153 in chromatin architecture and transcription by mediating CTCF and cohesin binding in mammalian cells. We propose that NUP153 links NPCs to chromatin architecture allowing developmental genes and IEGs that are poised to respond rapidly to developmental cues to be properly modulated.

## INTRODUCTION

Establishment of cell lineage specification, maintenance of cellular states and cellular responses to developmental cues rely on gene regulation and spatial genome organization during development ^1, 2, 3^. Emerging data point to highly coordinated activity between epigenetic mechanisms that involve nuclear architecture, chromatin structure and chromatin organization ^4, 5, 6, 7, 8^. However, our understanding on how nuclear architectural proteins are causally linked to chromatin organization and impact gene regulation have been limited underscoring the importance of defining the molecular determinants.

Nuclear architecture is in part organized by the nuclear lamina composed of lamin proteins and the nuclear pore complex (NPC). Nucleoporin proteins (Nups) are the building blocks of the NPC, which forms a ∼60-120 mega dalton (mDa) macromolecular channel at the nuclear envelope mediating nucleocytoplasmic trafficking of proteins and RNA molecules during key cellular processes such as cell signal transduction and cell growth (reviewed in ^9^). Beyond their role in nuclear transport, the NPC has been one of the nuclear structural sites of interest for its potential role in gene regulation by directly associating with genes (reviewed in ^10^). Studies in budding yeast and metazoans have shown that the NPC provides a scaffold for chromatin modifying complexes and transcription factors, and mediates chromatin organization. In metazoans, such compartmentalization supports nucleoporin-chromatin interactions that influence either transcriptional activation or silencing ^11, 12, 13, 14^. In yeast, the majority of the genes that position to the NPC are transcriptionally active. For example, inducible genes such as *GAL*, *INO1*, and *HXK1* relocalize from the nucleoplasm to the NPC upon transcription activation - a process that has been proposed to be critical for establishment of transcription memory ^15, 16, 17, 18, 19^. For several of these loci, NPC association facilitates chromatin looping between distal regulatory elements and promoters ^20, 21^. This mechanism has been proposed to be critical for expression and transcriptional memory of developmentally regulated ecdysone responsive genes in *Drosophila.* Upon activation, ecdysone responsive genes exhibit NUP98-mediated enhancer-promoter chromatin looping at the NPC ^22^. Notably, NUP98 has been shown to interact with several chromatin architectural proteins, including the CCCTC-binding factor, CTCF. These findings collectively suggest that Nups can facilitate chromatin structure in a direct manner by regulating transcription and in an indirect manner whereby Nup-mediated gene regulation relies on architectural proteins. Nevertheless, the functional relevance of Nup-architectural protein interactions in transcription regulation and chromatin structure is not well understood.

Chromatin architectural proteins, CTCF and the cohesin complex, facilitate interactions between *cis*-regulatory elements ^23, 24^. These interactions influence the formation and maintenance of long-range chromatin loops that underlie higher order chromatin organization ^25, 26, 27, 28^. Long-range loops of preferential chromatin interactions, referred to as “topologically associating domains” (TADs) are stable, conserved across the species, and exhibit dynamicity during development ^24, 29^. Importantly, TADs segregate into transcriptionally distinct sub-compartments ^30, 31^ and exhibit spatial positioning ^32^. Current models argue that lamina-chromatin interactions may provide sequestration of specific loci inside the peripheral heterochromatin and promote formation of a silent nuclear compartment ^33, 34^. Despite the close interaction between the nuclear lamina and the NPC, we still know very little on how NPC-chromatin interactions influence transcription and chromatin organization at the nuclear periphery.

In mammals, Nups show variable expression across different cell types and their chromatin binding has been attributed to cell-type specific gene expression programs (reviewed in ^10^). NUP153 is among the chromatin-binding Nups which have been proposed to impact transcription programs that mediate pluripotency and self-renewal of stem cells in mammalian cells ^35, 36, 37^. NUP153 chromatin binding sites have been detected at the promoters and across gene bodies ^35, 37^. Large proportion of NUP153 sites have also been detected at the intergenic sites containing enhancers that are linked to cell identity genes ^36^. Furthermore, the epigenetic and transcriptional state of NUP153 bindings sites show variability whereby NUP153 can associate with both transcriptionally active and silent regions of the genome ^35, 37^. Nevertheless, the molecular basis for how NUP153 association at the enhancers or promoters impact chromatin structure and transcription remain to be open questions.

Here, we directly tested the relationship between NUP153-chromatin interactions and gene regulation in pluripotent mouse ES cells. Towards elucidating NUP153-mediated mechanisms that control transcriptional silencing vs activation, we further utilized immediate early genes (IEGs) as model loci. We report that NUP153 interacts with chromatin architectural proteins, cohesin and CTCF, and mediates their binding at enhancers, transcription start sites (TSS) and TAD boundaries in mouse ES cells. NUP153 depletion results in differential gene expression that is most prevalent at bivalent genes ^8^. To determine the mechanism by which NUP153 regulates CTCF and cohesin function during gene expression, we utilized IEGs, including *Egr1*, *c-Fos* and *Jun* loci, which we identified as NUP153 targets in mouse ES cells. We took advantage of the fact that transcription at the IEG loci can be efficiently and transiently induced using growth hormones such as the epidermal growth factor (EGF) in HeLa cells ^38^. We found that NUP153 binding at the IEG *cis*-regulatory elements is critical for CTCF and cohesin binding and subsequent POL II pausing. We also found that this function of NUP153 is essential for efficient transcription initiation of IEGs. Notably, IEGs exhibit a NUP153-dependent peripheral positioning during the basal state and reposition even closer to the periphery during transcriptional activation. Our findings reveal that IEG-NUP153 contacts are essential for IEG transcription via establishment of a chromatin structure that is permissive for POL II pausing at the basal state. Collectively, we propose that NUP153 is a key regulator of chromatin structure by mediating binding of the chromatin architectural proteins, CTCF and cohesin, at *cis*-regulatory elements and TAD boundaries in mammalian cells. Through this function, NUP153 links NPCs to chromatin architecture allowing developmental genes and IEGs that are poised to respond rapidly to developmental cues to be properly modulated.

## RESULTS

### A proteomics screen identifies cohesin subunits as NUP153 interacting proteins

To understand the functional relevance of NUP153 in transcriptional regulation and chromatin structure, we utilized an unbiased proteomics screen using mouse NUP153 as bait in an affinity purification assay. We expressed FLAG-tagged mouse NUP153 (FLAG-mNUP153) in HEK293T cells and carried out immunoprecipitation (IP) followed by mass spectrometry (MS) (Figure 1A, S1A-B). We identified several known NUP153 interacting proteins including TPR ^39^, NXF1 ^40^, IPO5 ^41^, SENP1 ^42^, XPO-1, TNPO1, and RAN ^43^. In addition, IP-MS revealed that NUP153 interacts with several chromatin interacting proteins including the cohesin complex components, SMC1A, SMC3 and RAD21 (Figure 1A and data not shown).

**Figure 1:**
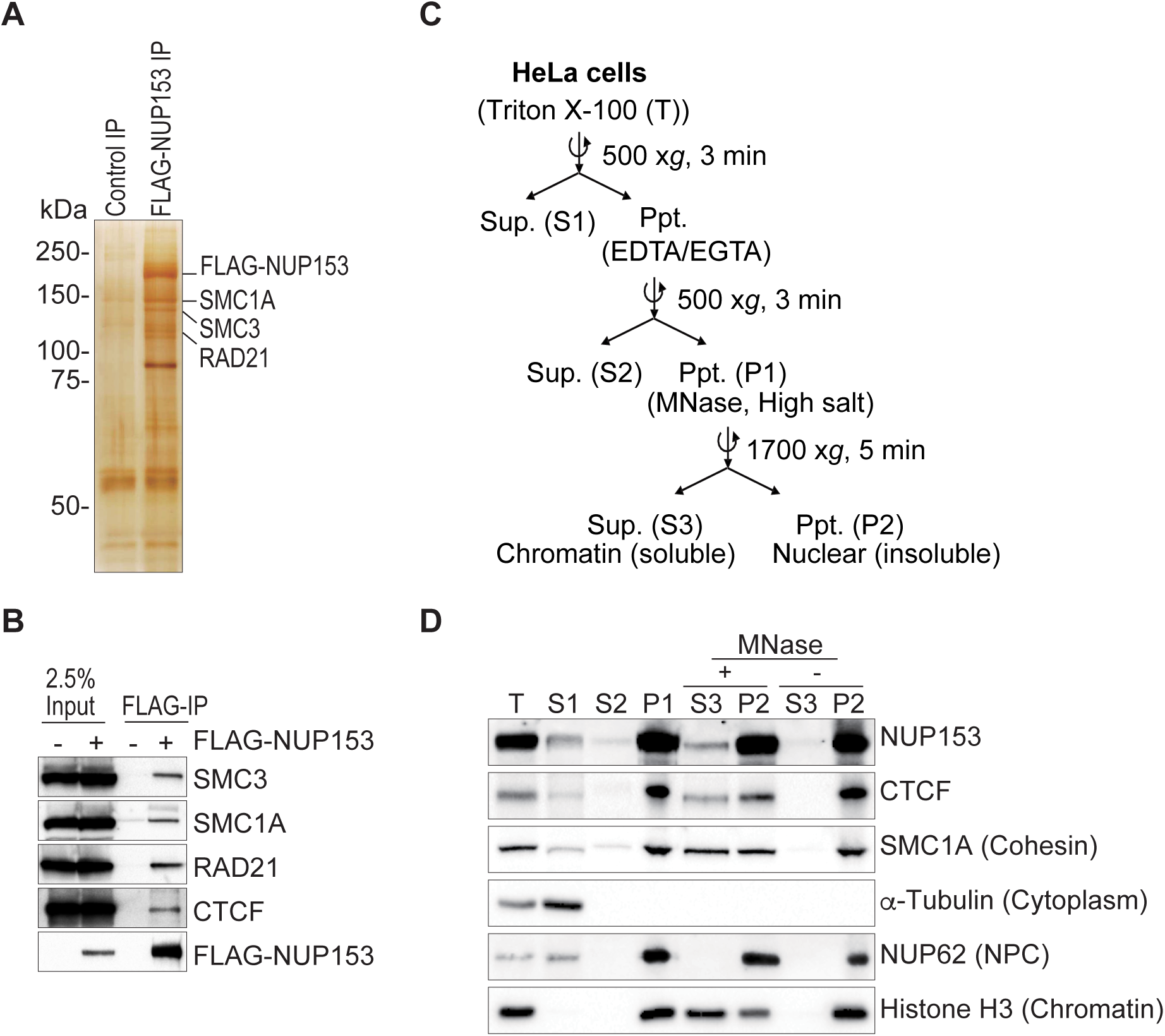
NUP153 interacts with CTCF and cohesin. (A) Silver stain showing proteins that IP with FLAG-NUP153. (B) Co-IP shows FLAG-NUP153 interaction with CTCF, and cohesin subunits, SMC3, SMC1A, RAD21. NUP153 was pulled down using anti-FLAG antibody (Ab). (C) Schematic showing steps of chromatin fractionation assay in HeLa cells. (D) NUP153 was detected in the nuclear insoluble fraction (P2) along with CTCF and cohesin. NUP153 detected in the chromatin-associated soluble fraction (S3) following micrococcal nuclease (MNase) treatment of P1 fraction. Ppt, precipitate; Sup, supernatant; Nucleoporin 62, NUP62; Loading controls: *α*-Tubulin (cytoplasm), Histone H3 (chromatin).

NUP153 has been mapped to enhancers and promoters in mammalian cells and has been implicated in transcription regulation ^35, 36, 37^. Nevertheless, whether NUP153 influences higher-order chromatin structure and how NUP153 impacts gene expression are not well understood. We, thus focused on the cohesin complex as it mediates higher-order chromatin organization, and regulates gene expression by facilitating and stabilizing enhancer-promoter interactions together with CTCF ^44, 45^. Cohesin binding sites show ∼70-80% overlap with CTCF chromatin interaction sites ^46^. It has also been recently shown that cohesin positioning to CTCF binding sites is influenced by both transcription and CTCF ^46, 47^. To investigate functional communication between NUP153, CTCF and cohesin, we performed FLAG-NUP153 IP followed by western blotting and determined NUP153 interaction with CTCF and cohesin subunits (Figure 1B).

To define the nuclear fraction at which NUP153 spatially interacts with CTCF and cohesin, we performed biochemical chromatin fractionation assay ^48^ using HeLa cells (Figure 1C). We successfully collected the core histone, Histone H3, in the nuclear fraction (P1). Treatment of the nuclear fraction (P1) with the micrococcal nuclease (MNase) resulted in elution of chromatin binding proteins into the soluble fraction (S3) in comparison to the insoluble nuclear fraction (P2) (Figure 1C-D). We detected the NPC component, NUP62, in the nuclear insoluble fraction (P2) in the absence or presence of MNase suggesting that the P2 fraction contains the intact nuclear membrane including the nuclear envelope and the NPC. Insoluble fraction has been also shown to contain several proteins, such as CTCF, that associate with the nuclear matrix ^49^. We detected NUP153 both in the nuclear insoluble (P2), and the soluble (S3) fractions that contain chromatin-binding proteins (Figure 1D). This data provides biochemical evidence supporting earlier cell biological reports that NUP153 associates with the nuclear membrane and is found as a soluble protein within the nucleoplasm ^50, 51, 52^. It also suggests that the NUP153-chromatin interactions might be established either at the nuclear periphery or in the nucleoplasm. Interestingly, similar to NUP153, we detected a proportion of CTCF and cohesin in the insoluble nuclear fraction (P2) even in the presence of MNase (Figure 1D). These findings argue that NUP153 may interact with CTCF and cohesin at the nuclear periphery, nuclear matrix or within the nucleoplasm.

### NUP153 enrichment at the *cis*-regulatory elements and TAD boundaries in pluripotent mouse ES cells

NUP153 mediates transcription regulation of developmental genes in mouse ES cells ^35^. Such function has been attributed to the transcriptional silencing role of NUP153 together with PRC1 complex. Nevertheless, only ∼10% of NUP153 binding sites overlap with PRC1 interaction sites explaining only a small proportion of NUP153-mediated gene regulation in pluripotent mouse ES cells. To study regulatory roles of NUP153 in chromatin structure and gene regulation, we generated female mouse ES cell lines (16.7) ^53^ that express NUP153 in fusion with *E. coli* DNA adenine methyltransferase (Dam) (Figure S1C-E) and mapped NUP153 chromatin interaction sites by DamID-Seq (Yildirim et al. unpublished). A Dam only expressing cell line was used to normalize DamID-Seq data for chromatin accessibility differences across the genome. The DamID method has been successfully used to map protein-chromatin interactions in mammalian cells ^54^. The method relies on the ability of *E. coli* Dam enzyme to catalyze adenine N^6^ methylation (6mA) within the GATC sequences of DNA sites, which show association with the fusion protein of interest ^55, 56^. We identified 73,018 high confidence NUP153 binding sites (greater than 2-fold enrichment over Dam-only control and FDR<0.05). We examined the distribution of NUP153-DamID Seq peaks across genetic elements. In agreement with earlier reports, ^35^ (Yildirim et al. unpublished), we detected 32.2% of the NUP153 DamID peaks at intergenic sites, 14.2% of peaks at promoters, and 53.5% of peaks across gene bodies in female mouse ES cells (Figure 2A).

**Figure 2:**
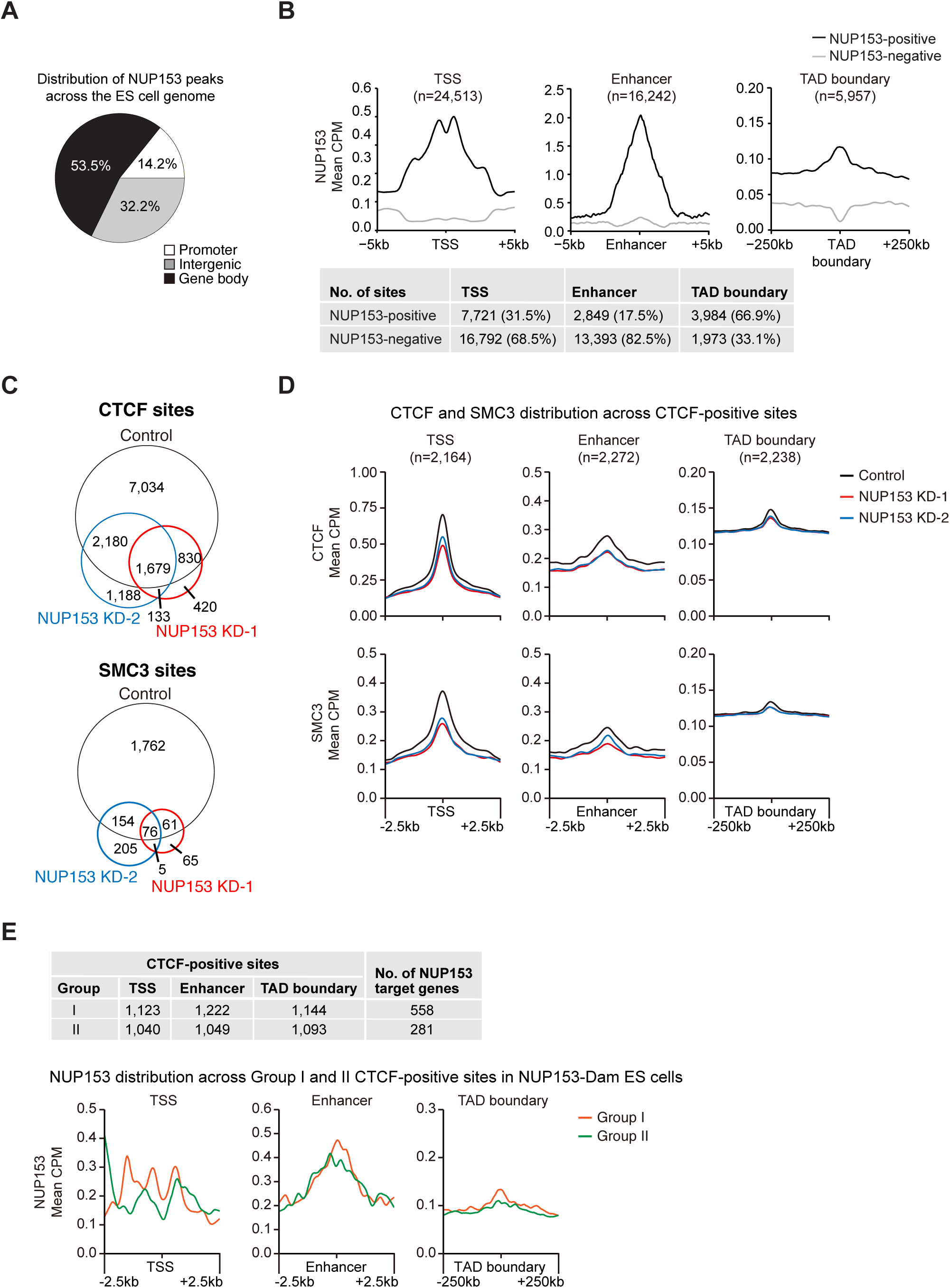
NUP153 mediates CTCF and cohesin binding at *cis*-regulatory elements and TAD boundaries in mouse ES cells. (A) Distribution of NUP153 peaks in mouse ES cells. Peaks are categorized as promoters (×2kb from TSS to +100 bp from TSS), gene body (+100bp from TSS to +1kb from transcription termination site (TTS)), intergenic sites (< −2kb from TSS and >+1kb from TTS). (B) Metagene profiles of mean NUP153 binding at NUP153-positive and NUP153-negative TSS and enhancer (+/− 5kb) and TAD boundaries (+/-250kb) (top). Number and percentage of NUP153 binding sites are presented as a table for the indicated genetic elements (bottom). (C) Genome-wide CTCF and SMC3 binding sites were compared in control and NUP153 deficient (KD-1, KD-2) mouse ES cells. (D) Metagene profiles showing mean CTCF and SMC3 binding across CTCF-positive TSS (n=2,164), enhancers (n=2,272) and TAD boundaries (n=2,238) in control and NUP153 deficient mouse ES cells. (E) Mean CTCF binding in control and NUP153 KD cells were compared and CTCF sites were grouped into two. Group I, contained CTCF sites that showed greater mean CTCF binding in control cells over NUP153 KD cells. Group II contained CTCF sites that showed equal or lesser mean CTCF binding in control cells over NUP153 KD cells. Number of CTCF-positive sites across TSS and enhancer (+/− 2.5kb) and TAD boundaries (+/− 250kb), and the number of NUP153 target genes that associate with each Group are shown as a Table (top) (See also Figure S3D). Metagene profiles showing mean NUP153 binding across CTCF-positive Group I and Group II TSS, enhancer (+/− 2.5kb) and TAD boundaries (+/-250Kb) (bottom).

We next examined NUP153 distribution across three genetic elements including the TSS, enhancers and TAD boundaries (Figure 2B). NUP153 binding was detected at the TSS (Figure S2A) and we identified 31.5% of TSS (7,721/24,513) to be NUP153-positive (Figure 2B). To investigate the transcriptional state of the NUP153 chromatin binding sites, we performed RNA-Seq in ES cells and determined transcriptionally active vs inactive TSS based on Fragments Per Kilobase of transcript per Million mapped reads (FPKM). TSS with FPKM>1 (n=8,768) were denoted as active and TSS with FPKM*≤*1 (n=11,861) were denoted as inactive. By utilizing previously published Histone 3 Lysine 4 trimethylation (H3K4me3) and H3K27me3 ChIP-Seq data sets ^57^, we validated transcriptional activity and silencing at these sites (Figure S2B). We found that NUP153 occupied both transcriptionally active and inactive TSS with a bias towards the active genes (Figure S2B).

To evaluate NUP153 binding across enhancers, we mapped enhancers (n=16,242) using previously published ChIP-Seq against enhancer specific histone marks, H3K4me1 ^58^, Histone 3 Lysine 27 acetylation (H3K27Ac) ^59^ and Chromatin Binding Protein (CBP)/P300 ^60^ (Figure S2C). We detected NUP153 enrichment at the enhancers (Figure S2C) and identified ∼17.5% NUP153-positive enhancers (2849/16242) (Figure 2B). Compared to NUP153-negative enhancers, NUP153-positive enhancers exhibited higher H3K4me1, H3K27Ac and CBP/P300 occupancy (Figure S2D). Distribution of NUP153 at the TSS and enhancers suggested that NUP153 may have a functional role in gene regulation.

Considering interaction of NUP153 with CTCF and cohesin complex, the third genomic region we focused on was the TAD boundaries which are characterized to be enriched for CTCF and cohesin (^61^ (reviewed in ^62^)). To identify TAD boundaries, we utilized the previously reported coordinates based on Hi-C data in mouse ES cells ^61^. We found that 66.9% of TAD boundaries (3,984/5,957) contained NUP153 binding (Figure 2B) suggesting that NUP153 may functionally cooperate with CTCF and/or SMC3 at the TAD boundaries during chromatin organization.

### NUP153 mediates CTCF and cohesin binding at *cis*-regulatory elements and TAD boundaries

CTCF and cohesin binding across the intergenic sites have been linked to their role in chromatin insulation or enhancer function during gene expression ^63, 64^. To determine the functional relevance of NUP153 interaction with CTCF and cohesin during gene regulation, we first mapped CTCF and cohesin binding sites by ChIP-Seq and correlated the data with NUP153 DamID-Seq to define co-occupied sites. In accordance with earlier reports ^64^, CTCF and SMC3 were enriched across TSS, enhancers (Figure S2A, S2C) and TAD boundaries (Figure 2D). We found that on average CTCF and cohesin binding sites were at ∼5 kb distance with respect to the nearest NUP153 binding sites (Figure S2E). Based on this criterion, we detected a robust co-localization whereby 48.9% of the CTCF and 44.4% of the SMC3 binding sites were co-occupied by NUP153. Out of the CTCF^+^/NUP153^+^ co-occupied sites, 29.9% associated with TSS, and 24.2% associated with enhancers. SMC3^+^/NUP153^+^ co-occupied sites presented a similar profile in that 23.9% of these sites associated with TSS, and 27.1% associated with enhancers. By examining sites that are occupied by all three factors (NUP153^+^/CTCF^+^/SMC3^+^), we found association of the sites with 10.4% of the TSS and 13.9% of the enhancers. These results pointed to a potential crosstalk between NUP153 and architectural proteins during regulation of gene expression and/or chromatin architecture.

To investigate the functional relevance of NUP153 in CTCF and cohesin distribution genome-wide and define its impact on gene regulation, we generated NUP153 knockdown (KD) ES cells by transducing cells with two different mouse NUP153-specific shRNA lentivirus. We confirmed ∼55-60% downregulation of NUP153 expression by real-time PCR (Figure S2F). Both control and NUP153 deficient cells showed typical pluripotent ES cell characteristics with their morphology and the presence of alkaline phosphatase activity ^65^, suggesting that NUP153 depletion did not interfere with the pluripotent state of ES cells (Figure S2G). By utilizing an oligo (dT)50-mer probe and performing RNA Flourescent *in situ* hybridization (FISH) ^66^, we further validated that the Poly(A)^+^ RNA export function of the NPCs was intact in NUP153 KD ES cells (Figure S2G).

We next evaluated how NUP153 depletion influenced distribution of CTCF and cohesin sites across different genetic elements (Figure S3A). In control ES cells, 22.8% of CTCF binding was detected at promoters, while 45% binding was detected across gene bodies and 32.2% of binding was detected at intergenic sites. In contrast, both NUP153 deficient ES cells exhibited a higher CTCF binding at promoters (30-34%) and lower gene body- (38-41%) and intergenic- (27-29%) specific CTCF binding. Similar distribution patterns were detected for SMC3 in control and NUP153 KD cells (Figure S3A). These results suggested that NUP153 may impact CTCF and cohesin binding across the genome. To this end, we compared number of CTCF and cohesin peaks between control and NUP153 KD ES cells (Figure 2C). We identified a significant loss of genome-wide CTCF (∼60%) and SMC3 (∼86%) binding in NUP153 deficient ES cells (Figure 2C).

Given that cohesin binding relies on CTCF ^46, 67^, we focused on CTCF binding sites and showed that NUP153 is enriched at the CTCF-positive TSS (n=2,164; p=0, hypergeometric test), enhancers (n=2,272; p=0, hypergeometric test) and TAD boundaries (n=2,238; p=8.66e-103, hypergeometric test) (Figure S3B). We next asked whether NUP153 regulates CTCF and/or cohesin binding selectively at a given genetic element and found that NUP153 depletion resulted in reduction in CTCF and cohesin binding across all three genetic elements (Figure 2D). We next asked how NUP153 binding influence CTCF distribution. To address this question, we calculated the mean CTCF binding in NUP153 deficient cells in comparison to control cells and grouped the CTCF binding sites into two. Group I, contained CTCF sites that showed greater mean CTCF binding in control cells over NUP153 KD cells, and Group II contained CTCF sites that showed equal or lesser mean CTCF binding in control cells over NUP153 KD cells (Figure 2E, S3C-E). Group I TSS sites constituted ∼10% (1,123/11,726) of the total CTCF binding sites and half of these sites (∼5%, 558/11,726) were NUP153 positive. Notably, metagene profiles across TSS, enhancer and TAD boundaries showed higher NUP153 binding at Group I sites over Group II sites (Figure 2E). This data suggested that the degree of NUP153 binding correlates with differential change in CTCF binding at each genetic element. Based on these findings, we concluded that NUP153 mediates CTCF and cohesin binding at TSS, enhancer and TAD-boundaries. This raises the possibility that NUP153 may be critical for enhancer-promoter functions or chromatin organization functions of CTCF and cohesin during gene expression.

### NUP153-mediated transcription regulation across the genome and at bivalent genes

Given the enriched association of NUP153 with gene regulatory elements and its influence on CTCF and cohesin binding at TSS and enhancers, we investigated the extent of transcriptional changes in NUP153 deficient ES cells. To this end, we performed RNA-Seq for control and two NUP153 KD ES cell lines. NUP153 depletion resulted in differential expression of 711 genes (fold change ≥1.5 and FDR<0.05) genome-wide (Figure 3A). Approximately 56% (398/711) of this gene set contained TSS- or gene body specific NUP153 binding sites. Compared to control ES cells, a majority (66.2%, 471/711) of the differentially regulated genes were upregulated in NUP153 KD ES cells. Gene ontology (GO) analyses has revealed that the upregulated genes were associated with pathways such as those that impact cell differentiation (e.g. *Fgf1*, *Fgf9*, *Dlk1*, *Bmp7, Hoxb13)*, cell proliferation (e.g. *Cdx2*, *Ntrk3*, *Ebb4)*, and transcription regulation (e.g. *Wnt7b*, *Gata3*, *Bcl11a*, *ApoB, Lhx1, Pou3f2*). In contrast, expression of genes that regulate biological processes such as extracellular matrix organization (e.g. *Fbln5*, *Comp*, *Ntn4*, *Dmp1*), response to mechanical stimulus (e.g. *Cav1*, *Cxcl12*, *Col3a1*), and skeletal muscle development (e.g. *Mef2c*, *Foxp2*, *Meox2*) were downregulated in NUP153 deficient ES cells.

**Figure 3:**
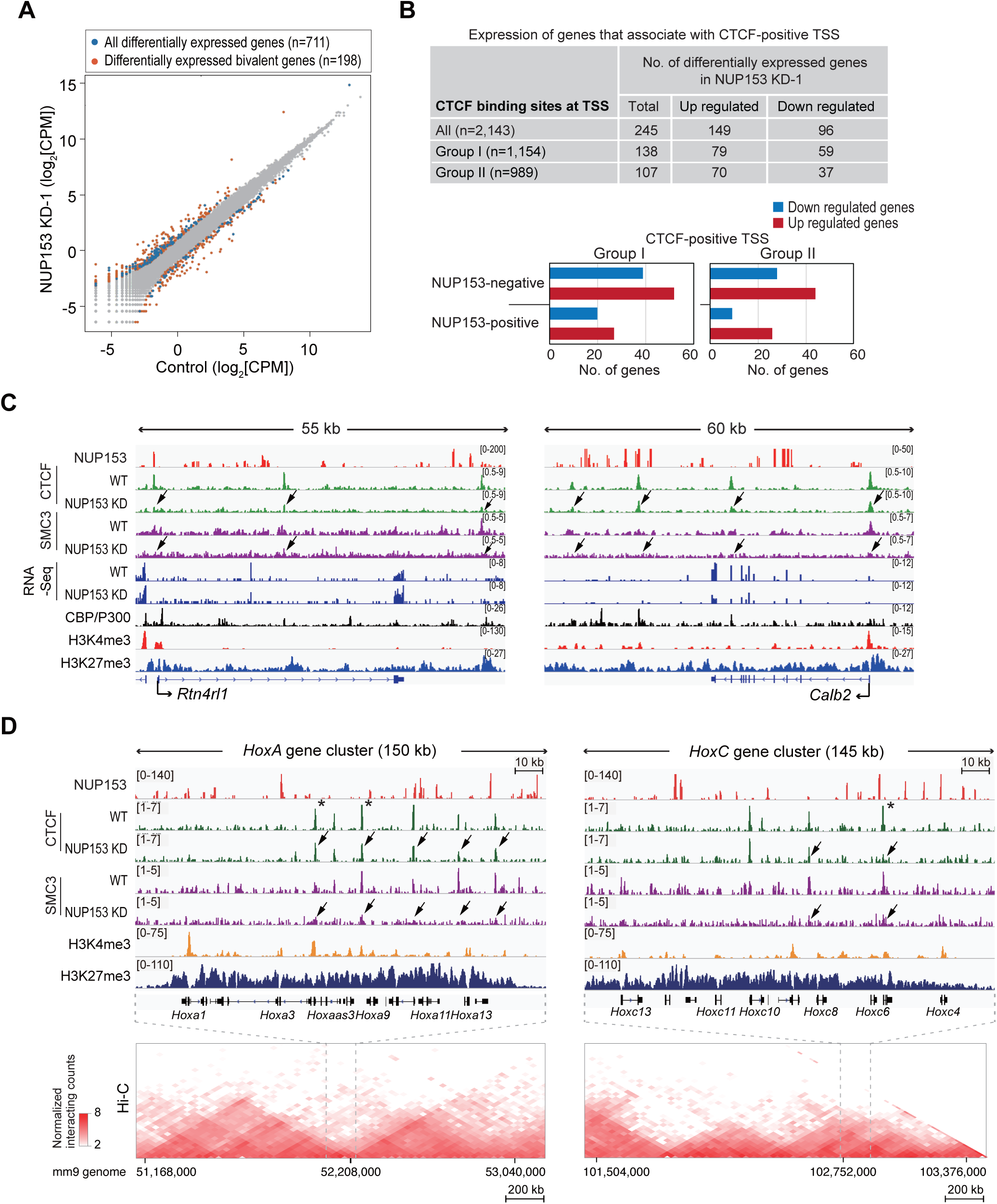
NUP153 influences transcription and binding of CTCF and cohesin at bivalent genes. (A) Scatter plot showing transcript expression levels in Log2[CPM] scale in control and NUP153 KD-1 ES cells. Blue points are all differentially expressed genes (n=711) and orange points are differentially expressed bivalent genes. (B) Table showing number of differentially expressed genes that associate with all, Group I and Group II CTCF-positive TSS (top). Plots showing number of differentially regulated NUP153-positive and NUP153-negative genes that associate with Group I and II CTCF-positive TSS (bottom). (C) NUP153 DamID-Seq, CTCF, cohesin, H3K4me3, and H3K27me3 ChIP-Seq, and transcripts (RNA-Seq) tracks are shown for two NUP153-positive Group I genes, *Rtn4rl1* (left panel) and *Calb2* (right panel) in control (WT) and NUP153 KD ES cells. *Rtn4rl1* shows transcriptional up regulation and *Calb2* shown transcriptional down regulation. (D) NUP153 DamID-Seq, CTCF, cohesin, H3K4me3, and H3K27me3 ChIP-Seq tracks are shown for a 145-150 kb region for *HoxA* and *HoxC* loci in control (WT) and NUP153 KD ES cells as indicated. Arrows point to regions where CTCF or SMC3 binding are altered in NUP153 KD ES cells. CTCF sites labeled with asterisk (*) point to CTCF sites that have been reported to regulate transcription at *Hox* loci by mediating formation of TADs ^71, 75^. The 2D heat map shows the interaction frequency in mouse ES cells ^61^. Hi-C data was aligned to the mm9 genome showing *HoxA* cluster residing in a TAD boundary and *HoxC* cluster in a TAD as published ^61^. H3K4me3 and H3K27me3 ^57^ and CBP/P300 ^60^ ChIP-Seq data were previously published. CPM, Counts per million.

We next investigated how NUP153-dependent changes in CTCF binding may impact transcription. We found that ∼34.4% (245/711) of the differentially regulated genes associated with CTCF-positive TSS. Majority of this gene set (∼61%) showed transcriptional upregulation (Figure 3B). Examining the biological function of these genes by GO analysis, we found that they associate with important cellular processes such as the cell migration (e.g. *Ptk2b*, *Tcaf2*, *Wnt11*), cell adhesion (e.g. *Alcam*, *App*, *Itga3*, *Itga8*, *PLCb1*), and cell differentiation (e.g. *Foxa3*, *Flnb*, *Zfp423*, *Tnk2*). Given that the CTCF-positive Group I sites showed drastic change in CTCF binding in NUP153 KD cells and they were enriched for NUP153 over Group II sites, we evaluated the number of differentially regulated genes between two groups. Group I sites associated with 19.4% (138/711) and Group II sites associated with 15% (107/711) of the differentially regulated genes in NUP153 KD-1 ES cells (Figure 3B). Notably, NUP153-positive Group I genes constituted ∼7% (47/711) of the differentially regulated genes. Genes that showed upregulation (57.4%, 27/47) versus downregulation (42.6%, 20/27) were almost equally distributed Representative tracks shown for NUP153-positive Group I genes *Rtn4rl1* and *Calb2* in Figure 3C present differential expression and altered CTCF and cohesin binding in NUP153 KD ES cells (Figure 3C). Collectively, these data suggested a regulatory role for NUP153 in global gene expression and for ∼7% (47/711) of the differentially expressed genes this function underlies NUP153-mediated CTCF binding at TSS.

Bivalent state of genes has been proposed to be critical for establishment and/or maintenance of the ES cell pluripotency transcription program ^1, 68, 69, 70^. A recent report by Mas *et al.* provided evidence that there is a causal relationship between the maintenance of bivalent state and chromatin organization in ES cells ^8^. We, thus examined the impact of NUP153 loss on global bivalent gene expression. We utilized the bivalent gene list (n=3,868) reported by Mas *et al.* in which bivalency was determined by the presence of Trithorax group protein, MLL2, in addition to the histone marks, H3K4me3 and H3K27me3, at the promoters ^8^. By cross-referencing the bivalent gene list with the list of genes that show differential regulation in NUP153 deficient ES cells, we identified 27.8% (198/711) of differentially regulated bivalent genes in NUP153 KD cells (p=1.21e-16, hypergeometric test) (Figure 3A). Of this gene set, 32.8% (65/198) were NUP153 target genes and ∼10% (20/198) associated with NUP153-positive Group I TSS sites. Even though this data supported a key role for NUP153 in regulation of transcription at bivalent genes, it suggested that expression of only a small proportion of bivalent genes is mediated through NUP153-mediated CTCF binding at TSS.

Our analyses on the bivalent genes revealed that several *Hox* genes, which are known to present bivalent state in ES cells ^1, 71^, are NUP153 targets. *Hox* loci exhibit a tightly controlled genomic organization that relies on TADs with enriched CTCF binding ^61, 72^. This organization has functional relevance during developmental expression of *Hox* genes ^73, 74^. As presented in the representative tracks shown for the *HoxA* and *HoxC* clusters, we found that NUP153 depletion resulted in altered CTCF and/or cohesin binding at specific *Hox* genes (Figure 3D, arrows). Importantly, three of these CTCF-binding sites (Figure 3D, asterisks) have been reported to be critical in facilitating formation of TADs and providing an insulator function during developmental regulation of *Hox* gene transcription in mouse ^71, 75^. Based on our findings, we propose that NUP153 may contribute to the higher-order chromatin organization by regulating CTCF and cohesin positioning at specific developmental genes and mediate their gene expression.

### NUP153-mediated POL II recruitment during the IEG paused state is critical for timely IEG transcription

To determine the mechanism by which NUP153 regulates CTCF and cohesin function during gene expression, we utilized IEGs including *Egr1*, *c-Fos* and *Jun* loci which we identified to be NUP153 targets in mouse ES cells (Figure S4). It is well established that transcription at the IEGs is regulated by a proximally paused-POL II release mechanism ^76, 77, 78^. This mechanism results in POL II occupancy at the promoter-proximal regions (20-50 bp downstream of TSS), allowing for rapid and transient responsiveness of the IEGs to stimuli such as growth hormones ^38^.

By examining transcription and chromatin structure across the IEG loci, we determined that the TSS and distal regulatory elements of IEGs were occupied by CTCF and cohesin (Figure S4). During the preparation of this manuscript, it was shown that IEG locus, *EGR1,* forms CTCF-mediated higher order chromatin structure which impacts *EGR1* transcription in HeLa cells ^79^. Based on these characteristics, we envisioned that the IEG loci would provide a powerful *in vivo* model to examine mechanisms of NUP153-dependent gene expression, and provide a mechanistic understanding for the interplay between NUP153 and architectural proteins.

Earlier studies have revealed that IEG transcription kinetics showed variability in ES cells and thus could not be stably measured in this cell system ^80^. To test the function of NUP153 in transcription regulation directly, we thus utilized HeLa cells. In these cells, IEG transcription can be reduced to a silent state by serum starvation and transcription initiation can be reproducibly induced within 15 minutes upon EGF treatment ^38^. We generated NUP153 KD HeLa cells by transducing cells with NUP153-specific shRNA lentivirus and detected ∼60-80% reduction in NUP153 expression in comparison to control cells by western blotting (Figure 4A). We validated that NUP153 knockdown did not alter the nucleocytoplasmic trafficking at the NPCs by quantitating dexamethasone (Dex) responsive GFP-tagged glucocorticoid receptor (GR) nuclear import and export ^81^ (Figure S5A and S5B). As in mouse ES cells, NUP153 KD HeLa cells did not present any defects in Poly(A)^+^ RNA export which was evaluated by oligo(dT)50-Cy3 RNA FISH in control and NUP153 KD HeLa cells (Figure S5C).

**Figure 4:**
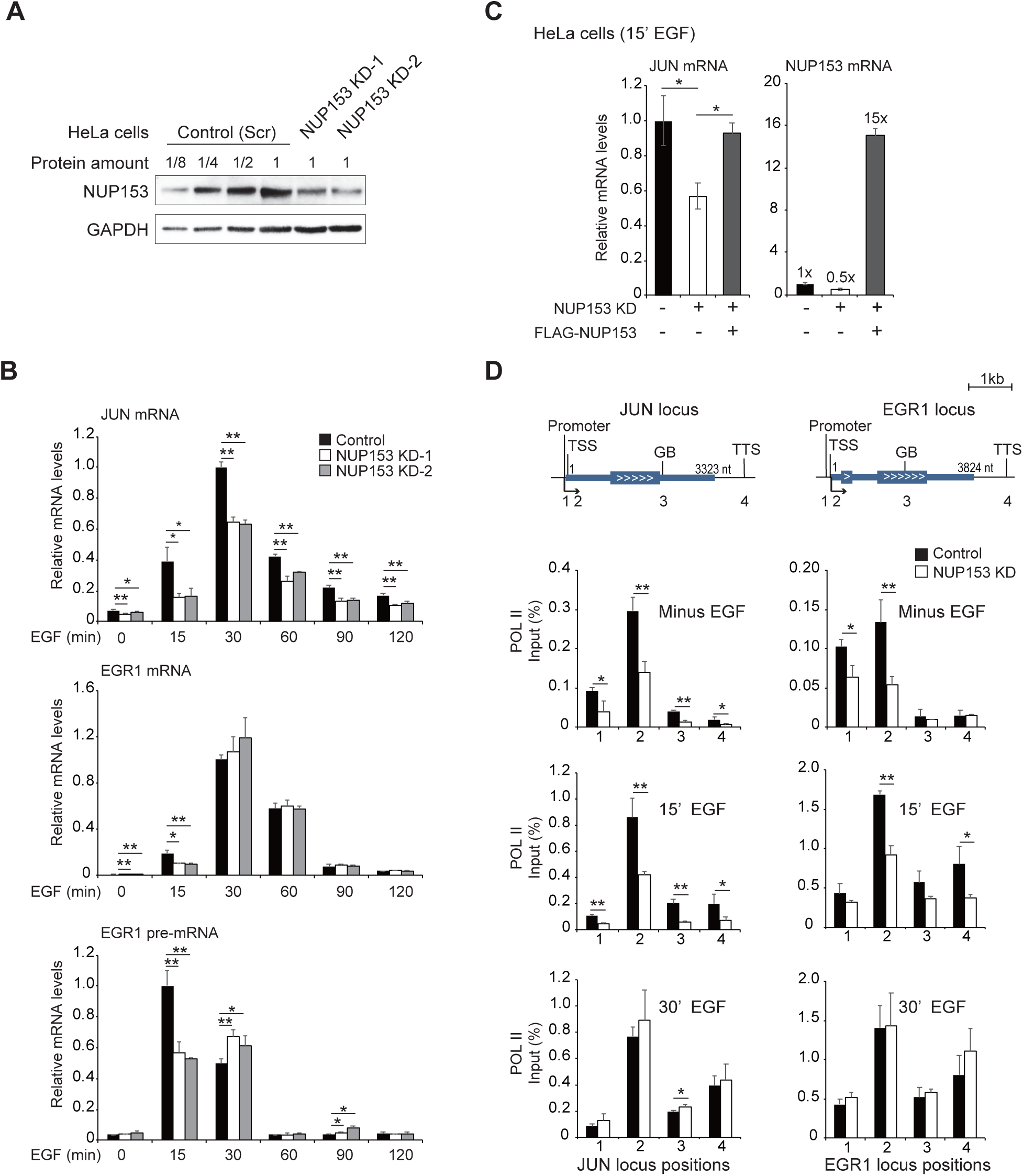
NUP153 controls POL II recruitment to IEG promoters and impact IEG transcription initiation. (A) Western blot showing NUP153 protein levels in HeLa cells transduced with control or NUP153 shRNA (KD-1 and KD-2) virus. (B) Real-time RT-PCR showing relative IEG mRNA and IEG nascent mRNA levels in control and NUP153 KD HeLa cells in a time course dependent manner. (C) Real-time RT-PCR showing relative IEG mRNA levels in control, NUP153 KD and NUP153 KD cells that express FLAG-NUP153 HeLa cells upon 15 min EGF treatment. *GAPDH* was used to normalize mRNA levels. (D) POL II binding across the IEGs, *JUN* and *EGR1* loci, was mapped by POL II ChIP in control and NUP153 KD HeLa cells at the paused state (minus EGF) and upon transcription induction 15 min and 30 min of EGF treatment. POL II occupancy at various IEG genetic elements (promoter, TSS, gene body (GB), and transcription termination site (TTS)) was measured using real-time PCR primers (see Table S1) as denoted in the schematics showing *EGR1* and *JUN* genes. Data shown are percent (%) of input at each genetic element. Values are mean ± standard deviation. Student’s t-test was applied to calculate significance. \**p*<0.05; \*\**p*<0.01. Experiments were repeated more than 3 times. Nt, nucleotide.

To further evaluate NUP153-dependent changes in IEG transcription, we utilized several IEG loci including *EGR1*, *JUN*, *c-FOS* and assessed the effects of NUP153 knockdown on IEG mRNA induction in response to EGF treatment in a time course dependent manner. Real-time PCR analyses have shown that transcription at all the IEGs examined was efficiently and transiently induced by EGF treatment at 15 min and the IEGs became transcriptionally silent by 120 min in control cells (Figure 4B, S6A and see Table S1 for list of primer sets used for real-time RT-PCR). Knockdown of NUP153 reproducibly led to a significant reduction in IEG mRNA and pre-mRNA levels upon 15 min of EGF treatment when compared to control cells (Figure 4B, S6A). We also detected a significant increase in the *EGR1* and *c-FOS* pre-mRNA levels upon 30 min EGF treatment (Figure 4B and S6A). These results argue that the suppression of IEG transcription during the initiation step leads to a delay in transcription or triggers a passive induction of negative feedback of IEG transcription ^38^. These transcriptional changes at the IEG were NUP153 specific, as expression of FLAG-NUP153 in NUP153 deficient HeLa cells led to recovery of transcription initiation (Figure 4C). These data collectively indicated that NUP153 acts as an activator of IEG transcription initiation.

Given that IEG transcription is mediated by the POL II pause-release mechanism ^76, 77, 78^, we reasoned that NUP153 may control POL II occupancy pre- and/or post-transcription induction. To investigate, we performed POL II ChIP and quantitatively measured POL II occupancy at the TSS and across gene bodies (GB) of *JUN* and *EGR1* using specific primer sets (Figure 4D, and see Table S1 for primers used for ChIP real-time PCR). At the paused state (minus EGF), NUP153 knockdown led to significant reduction in the paused POL II amounts at the IEG TSS. Upon 15 min of EGF induction, POL II occupancy at the TSS and across the gene body of IEGs was also significantly lower in NUP153 KD HeLa cells. In contrast, POL II binding across the IEGs was comparable between NUP153 KD and control cells at 30 min of EGF induction. These results were in line with the real-time PCR data showing that NUP153 is critical for timely IEG transcription initiation (Figure 4B). We concluded that NUP153 regulates IEG transcription initiation by controlling POL II occupancy across the TSS of IEGs at the paused state.

### CTCF and cohesin binding at *cis*-regulatory elements of IEGs relies on NUP153

In mouse ES cells, NUP153 depletion led to significant reduction in CTCF and cohesin binding at the TSS and enhancers coupled with differential changes in transcription (Figure 2, 3). Here, we took advantage of the inducible IEG loci to test the functional relationship between NUP153, and CTCF and cohesin during the paused state and transcription in a time course dependent manner. We utilized HeLa cell specific ENCODE ChIP-Seq data sets ^82^ (see Methods for details on the ENCODE datasets) and examined *EGR1* and *JUN* loci-specific POL II occupancy along with chromatin structure by mapping CTCF and cohesin subunit, RAD21, enhancer-specific marks (H3K27Ac, H3K4me1, and CBP/P300), and H3K4me3 which positively correlates with transcription activation (Figure S6B). Based on these maps, we designed primer sets to determine NUP153 kinetics at the IEG and evaluated NUP153-dependent changes in CTCF, cohesin occupancy across the predicted distal regulatory elements (enhancers) and IEG genetic elements including the TSS, promoter, GB and transcription termination sites (TTS) (Table S1).

NUP153 ChIP-Seq revealed that at the paused (minus EGF) state, NUP153 associates with the *EGR1* and *JUN* enhancers (site 7 for the *JUN* locus, sites 2 and 3 for the *EGR1* locus), TSS and TTS (Figure 5 (left panel)). Furthermore, we found that NUP153 binding spreads across the loci in a transcription dependent manner (Figure 5 (right panel)). These data suggested that the dynamics of NUP153 binding is tightly coupled to the transcriptional state of IEGs. Notably, similar to NUP153, CTCF and cohesin were also enriched around *EGR1* and *JUN* enhancer sites at the paused state and both proteins dynamically dissociated from these sites upon transcriptional activation with EGF. Furthermore, enhancer-specific binding of both proteins was dependent on NUP153. This is because CTCF and cohesin occupancy on IEG enhancers were significantly reduced in NUP153 KD cells compared to the control cells at the paused state (minus EGF), but were comparable between NUP153 KD and control cells in the transcriptionally activate state (+EGF) (Figure 5). A recent report provided evidence that EGR1 transcription relies on CTCF-mediated higher order chromatin ^79^. In the light of our findings, it is likely that NUP153 influences IEG chromatin organization by mediating CTCF and cohesin binding during the paused state.

**Figure 5:**
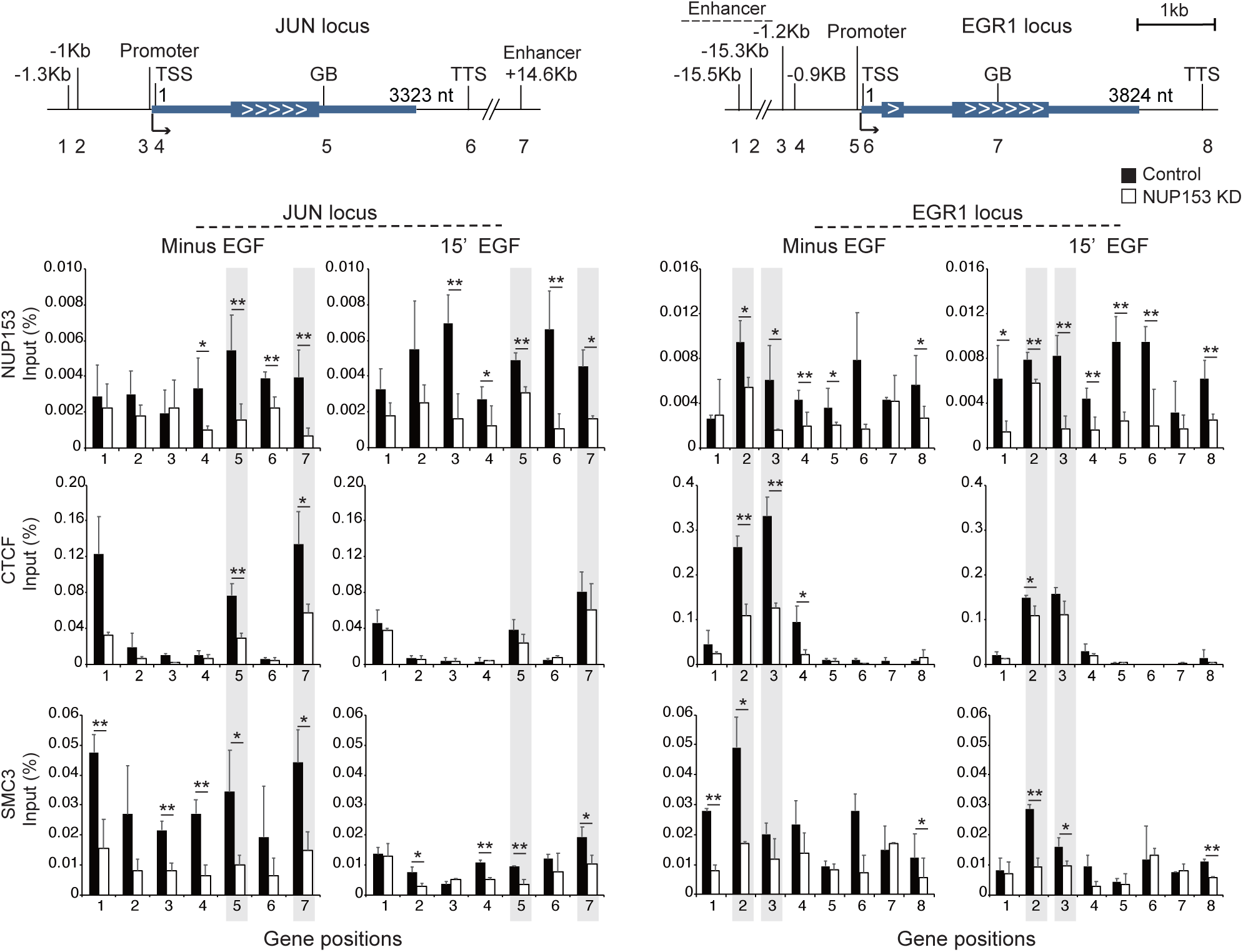
NUP153 is critical for CTCF and cohesin binding at the IEG *cis*-regulatory sites and across the IEG loci at the paused state. NUP153, CTCF and SMC3 occupancy across the JUN (left) and EGR1 (right) genetic elements were examined by ChIP real-time PCR at the paused state (minus EGF) and upon transcription induction (15 min EGF) in control and NUP153 KD HeLa cells. Position of PCR primers are denoted as numbers in the schematics showing EGR1 and JUN genes (See Table S1 for primer sequences). Data shown are percent (%) of input at each genetic element. Values are mean ± standard deviation. Student’s t-test was applied to calculate significance. \**p*<0.05; \*\**p*<0.01. Experiments were repeated more than 3 times. Nt, nucleotide.

### Co-regulatory function of NUP153 and CTCF during IEG paused state

Based on these results, we hypothesized that NUP153-mediated CTCF and cohesin binding to the IEG enhancer sites might be necessary for POL II occupancy at the proximal-promoter sites during the IEG paused state. Given that cohesin distribution depends on CTCF binding and POL II elongation ^47, 83, 84^, we focused on the functional relationship between NUP153 and CTCF. We generated CTCF knockdown HeLa cells by using shRNA against CTCF (Figure 6A) and measured impact of CTCF downregulation on transcription and POL II occupancy at the IEG loci in a time course dependent manner. Similar to the phenotype of NUP153 KD HeLa cells (Figure 4B), depletion of CTCF also resulted in significant reduction in IEG transcription initiation (Figure 6B). We also detected significant reduction in POL II occupancy at the promoter and TSS during the IEG paused state (minus EGF) (Figure 6C). Importantly, upon targeting both NUP153 and CTCF by shRNA (NUP153/CTCF KD), we did not detect an additive effect in downregulation of IEG transcription (Figure 6D) suggesting that NUP153 and CTCF mediate IEG transcription through the same regulatory mechanism.

**Figure 6:**
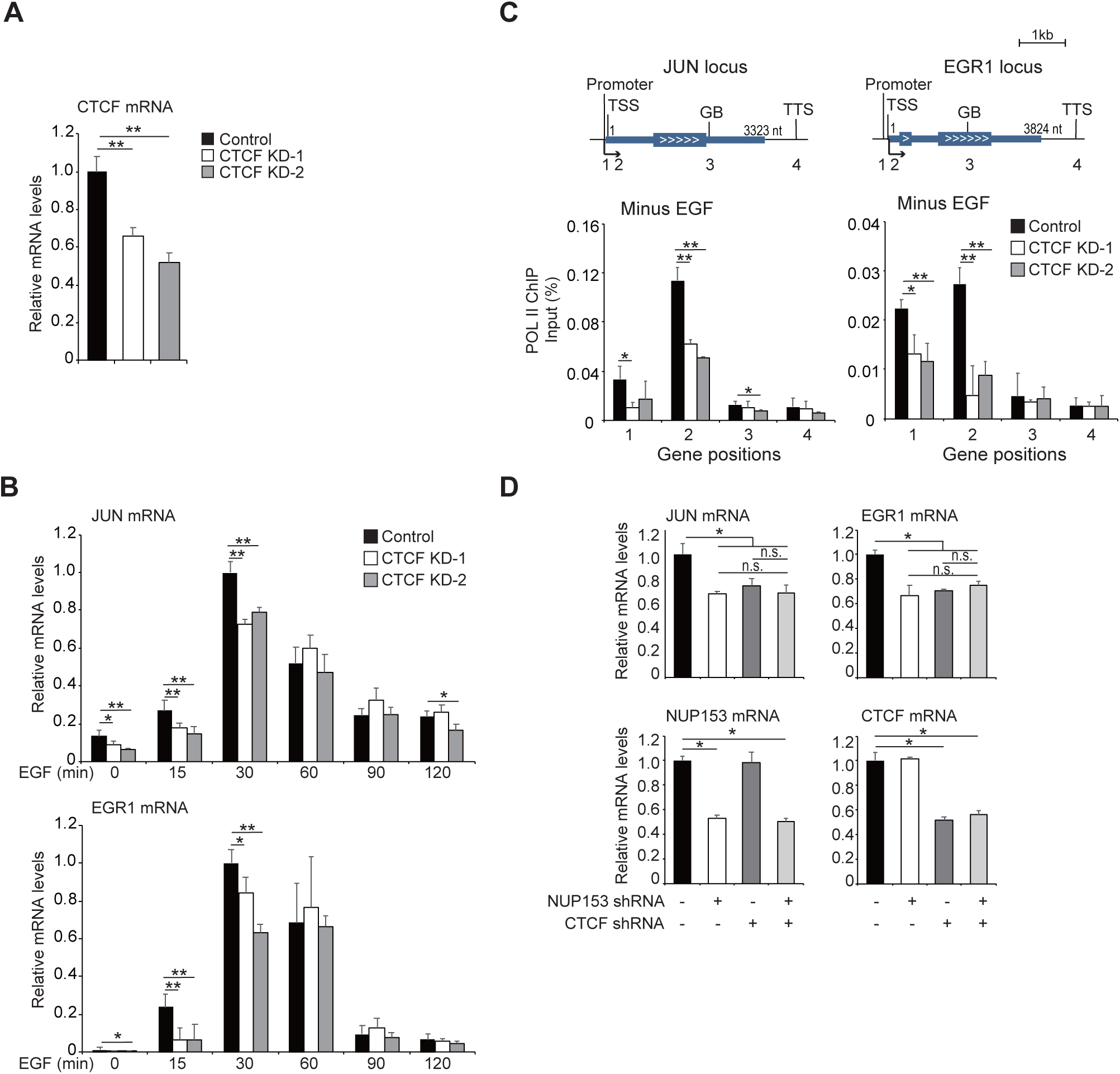
NUP153 and CTCF co-regulate POL II recruitment at the IEG paused state. (A) Real-time RT-PCR showing relative CTCF mRNA levels in HeLa cells transfected with control or CTCF shRNA expression vectors (KD-1 and KD-2). (B) Real-time RT-PCR showing relative *JUN* and *EGR1* mRNA levels in control or CTCF KD HeLa cells. (C) POL II binding across the IEGs, *JUN* and *EGR1* loci, was mapped by POL II ChIP in control and CTCF KD HeLa cells at the paused state (minus EGF). POL II occupancy at the indicated genetic elements was measured using real-time PCR primers as denoted in the schematics showing *EGR1* and *JUN* genes (Table S1). Data shown are percent (%) of input at each genetic element. (D) Real-time RT-PCR showing relative *EGR1*, *JUN*, *NUP153* and *CTCF* mRNA levels were measured in NUP153 KD, CTCF KD and in CTCF/NUP153 KD HeLa cells. Values are mean ± standard deviation. Relative mRNA levels were normalized using *GAPDH* mRNA levels. Student’s t-test was applied to calculate significance. \**p*<0.05; \*\**p*<0.01. Experiments were repeated more than 3 times. Nt, nucleotide.

### NUP153-dependent spatial positioning of IEGs during transcription regulation

Here, we investigated the spatial organization at the *c-FOS* locus and its dependency on NUP153 during the paused and transcriptionally active state in a time course dependent manner. To this end, we examined the sub-nuclear position of *c-FOS* DNA with respect to the nuclear periphery in control and NUP153 KD HeLa cells. We used a *c-FOS* gene containing Bacterial Artificial Chromosome (BAC) clone (RP11-293M10) as a DNA probe and performed DNA FISH in combination with immunofluorescence using an anti-Lamin B1 antibody to label the nuclear periphery (Figure 7A). We determined that the HeLa cells contained three *c-FOS* alleles. We utilized the microscopy images to measure the distance between each *c-FOS* locus and the nuclear periphery by Fiji software (see Methods for details on the calculation of normalized distance (ND) based on cell area). Analysis of cumulative frequency graphs has revealed that *c-FOS* locus is closely positioned (ND*≤*0.12) to the nuclear periphery in ∼30% of the control cells at the paused state (minus EGF) and that the loci moved even closer to the periphery (ND*≤*0.10) upon transcription induction (+EGF). In contrast, NUP153 deficiency led to positioning of the locus almost similar to the paused state in control cells whereby the locus remained distal to the periphery independent of the transcriptional state (Figure 7B and S7). These results argue that NUP153-dependent NPC positioning of IEG is critical during transcription regulation. These data also support our biochemical and genome-wide analyses showing that NUP153-mediates spatial positioning of CTCF and cohesin to the NPC during IEG transcription.

**Figure 7:**
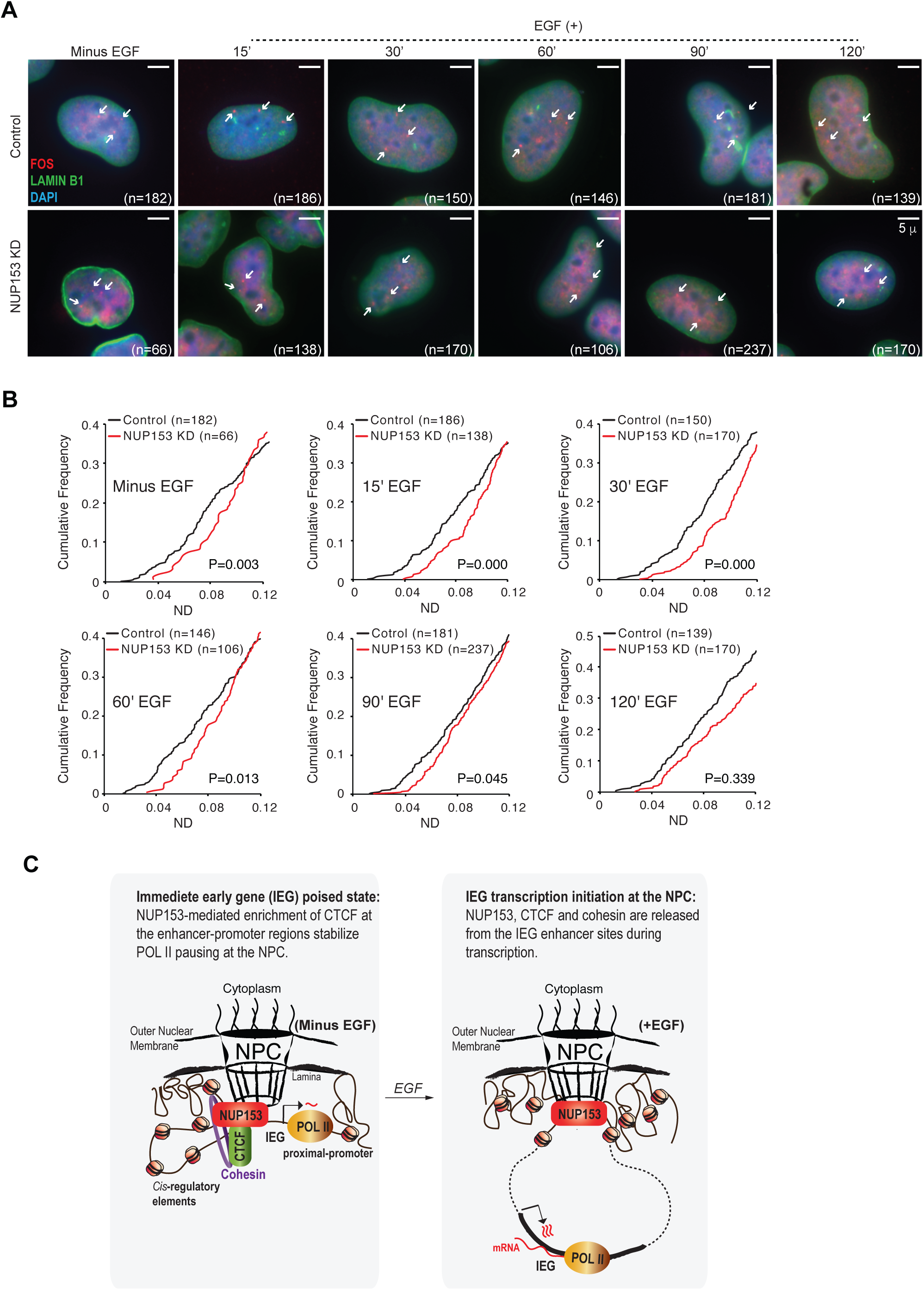
IEGs associate with the NPC in NUP153-dependent manner during paused state and transcription initiation. (A) Immunostaining of Lamin B1 and *c-FOS* DNA FISH in control and NUP153 KD HeLa cells are shown at the indicated time points. Cell numbers are as indicated. HeLa cells contain three *c-FOS* alleles (white arrows). Scale bar, 5μm. (B) Cumulative frequency graphs showing distribution of the *c-FOS* locus distance to nuclear periphery in control and NUP153 KD HeLa cells at the indicated time points. Cumulative frequencies at a normalized distance (ND) of 0.0-0.12 are shown. See Figure S7 for distribution of loci in all cells that were analyzed (ND of 0.0-0.5). ND= c-FOS locus to periphery distance/cell diameter (d), where d=(2xnuclear area/*π*)^0.5^. \**p*<0.05; \*\*\**p*<0.001; Kolmogorov–Smirnov (KS)-test was applied to calculate significance. Experiments were repeated twice. (C) Working model showing NUP153-mediated chromatin structure and transcription regulation at the IEG locus.

## DISCUSSION

In this study, we aimed to provide a mechanistic understanding on how NUP153 mediates chromatin structure and influences transcription. Through an unbiased proteomics approach, we identified NUP153 association with chromatin architectural proteins, CTCF and cohesin, and revealed that NUP153 is a critical regulator of chromatin structure and transcription by affecting CTCF and cohesin binding across enhancers and promoters in mammalian cells. Specifically, we determined that NUP153 depletion altered CTCF and cohesin binding in mouse ES cells at the *cis*-regulatory elements and TAD boundaries. Examination of the number of CTCF or SMC3 differential binding sites between the two different NUP153 KD mouse ES cell lines showed that they vary (Figure 2B). Nevertheless, CTCF and cohesin distribution patterns across different genetic elements showed similar pattern in different NUP153 KD cells (Figure S3A). These data argue that NUP153 facilitates CTCF and cohesin binding to their putative binding sites rather than recruiting them.

NUP153 is one of the mobile Nups that associates with the NPC basket or is found within the nucleoplasm ^50, 52^. Our data suggest that the co-regulatory function of NUP153 and architectural proteins likely occurs around the NPC. Even though we cannot exclude the fact that nucleoplasmic or nuclear matrix association is not possible, two of our results support NPC specific association. First, we have provided evidence that NUP153, CTCF and cohesin are detected in the nuclear insoluble fraction that contains the nuclear envelope and the nuclear matrix (Figure 1D). Second, we have shown that the IEG loci, which we show to rely on cooperation between NUP153 and architectural proteins for efficient transcription, present a close spatial positioning to the periphery, which is distorted upon NUP153 depletion (Figure 7, S7). There is already evidence in yeast showing that the inducible genes such as the *HXK1* facilitate chromatin looping between distal regulatory elements and promoters at the NPC ^20, 21^. A similar mechanism has been also proposed to be critical for NUP98-dependent regulation of ecdysone responsive genes during *Drosophila* development ^22^. Thus, use of mouse models or *in vitro* cell differentiation models will be necessary to further determine the regulatory role of NUP153 in chromatin organization and examine dependency of these processes to the subnuclear distribution of NUP153. These studies would provide critical insights on the significance of NUP153 in cell type-specific genome function and transcription regulation.

Regulation of bivalent gene expression has been proposed to be necessary for maintenance of ES cell pluripotency ^1, 8^. Bivalent state is established through the simultaneous catalytic activity of MLL and Polycomb Repressive Complex 2 (PRC2). Recent work in pluripotent ES cells suggests that MLL2 deficiency results in increased Polycomb binding coupled with loss of chromatin accessibility across the promoters and alterations in long-range chromatin interactions ^8^. These results suggest that maintenance of bivalent state might be causally linked to higher order chromatin organization during regulation of developmental gene expression. In this study, we provided evidence that NUP153 is enriched over enhancers, and TAD boundaries and is critical for CTCF and cohesin binding at these sites (Figure 2D). Notably, previous reports have shown that NUP153 occupancy at enhancers associates with cell-type specificity in human cells ^36^ and NUP153 depletion in adult stem cells impacts chromatin accessibility ^37^. Furthermore, similar to NUP153, CTCF exhibits variable binding patterns in different cell types and impacts cell type-specific transcription ^85, 86, 87, 88^. It is thus plausible to speculate that NUP153 might cooperate with CTCF and/or cohesin in higher order chromatin organization during the establishment of bivalent state and subsequently influence cell-type specific gene expression programs. Utilizing pluripotent ES cells and ES cell differentiation as systems to investigate the role of NUP153 in bivalency are thus interests for future studies.

CTCF has been well-characterized as a chromatin insulator ^89^ and Hi-C based chromatin studies revealed that both CTCF and cohesin mediate insulation of TADs ^23, 24^. Interestingly, CTCF depletion doesn’t result in disappearance of TADs that associate with active or inactive genome compartments, but depletion of cohesin eliminates formation of loop structures across the genome ^23, 24^. These results point to a hierarchical control of higher order chromatin organization through a functional cooperation between CTCF and cohesin. Chromatin compartments can also be established based on specific chromatin interactions with the lamina or the NPC ^32, 62, 90^. For example, studies in yeast already provide evidence that yeast NUP153 homologue Nup2 acts as an insulator at the nuclear basket ^91^. In mammalian cells, NUP153 has been associated with establishment of heterochromatin domains in interphase cells ^92^. Furthermore, NUP153 have been implicated compartmentalization in transcription factors at the NPC in response to activation of signal transduction pathways during cellular senescence, cell migration and cell proliferation ^89, 92, 93, 94^. Our results showing a functional cooperation between NUP153 and architectural proteins suggests that mammalian NUP153 may have a role in multistep organization and/or insulation of site-specific higher-order chromatin around the NPCs. NUP153 interaction with architectural proteins at the enhancers and/or TAD boundaries may promote formation of a chromatin compartment at which transcription factors downstream of specific signal transduction pathways associate with target loci- a compartment that can provide spatial and temporal organization of gene expression in response to cellular cues. Based on our findings, *Hox* loci and IEGs are among the NUP153 targets which may be subjected to such regulation. Collectively, our findings fill a gap in the field by providing evidence that NUP153 acts as a nuclear architectural component that associates with CTCF and cohesin mediating their binding across TSS, distal regulatory elements and TAD boundaries. Such function can be attributed to its potential role either in higher-order chromatin organization and/or transcription regulation. We have determined that transcription of human IEG loci and a small subset (∼5%) of the mouse ES cell genes rely on NUP153-mediated CTCF and/or cohesin binding at TSS. Our results were in accordance with earlier findings showing that only ∼10% of all TSS bound CTCF associated with promoter activity ^23^. Thus, future studies focusing on the role of NUP153 in chromatin structure and chromatin organization are critical.

Several genome-wide studies have shown that paused POL II distribution shows positive correlation with CTCF and cohesin binding across metazoan genomes ^95, 96, 97^. CTCF is thought to induce POL II pausing by creating “roadblocks” on the DNA template obstructing transcription elongation ^83^. In this study, we provide new evidence that NUP153 cooperates with CTCF in regulation of POL II occupancy at the IEG loci during paused state. Specifically, CTCF depletion results in altered POL II recruitment at the IEG loci during the paused state- a phenotype that mimics NUP153 depletion (Figure 6C). Knockdown of both CTCF and NUP153 did not result in an additive effect on POL II occupancy (Figure 6D) arguing that NUP153 and CTCF mediate IEG transcription through the same regulatory mechanism. During the preparation of this manuscript, two recent reports have supported our findings by showing that CTCF-mediated chromatin organization impacts IEG transcription ^79, 98^. Based on our data, we propose that NUP153 interacts with CTCF and mediates its binding at the *cis*-regulatory elements which subsequently leads to cohesin recruitment and chromatin looping between gene regulatory elements and/or TADs at the NPC. This state is essential for the establishment of a poised chromatin environment at which efficient transcription initiation can be rapidly induced through a POL II pause-release mechanism in response to stimuli (Figure 7C). NUP153-dependent localization to the NPC might thus provide an advantageous spatial position to genes that are poised to respond rapidly to developmental cues during ES cell pluripotency and/or differentiation. Time course dependent analyses of NUP153 distribution along the IEGs also allowed us to examine NUP153 dynamics during transcription. We determined that NUP153 exhibits a wide distribution across the promoter and the gene bodies of IEGs during transcriptional activation (Figure 5) suggesting that there might be a tight functional correlation between NUP153 and POL II activity during transcription. Chromatin sites that are engaged with stalled or active POL II might therefore allow for the differential NUP153 binding and can provide its selectivity towards transcriptionally silent or active chromatin domains, respectively.

We found that CTCF and cohesin binding sites were on average ∼5 kb distance from the nearest NUP153 binding sites (Figure S2E). Despite this fact, we detected a robust reduction in CTCF and cohesin binding at the *cis*-regulatory elements, suggesting that NUP153 may influence CTCF and cohesin binding directly or indirectly. One possible mechanism through which NUP153 can influence binding of architectural proteins is through the scaffold feature of NPCs. In various organisms, NPC acts as a scaffold at which specific genes associate with chromatin regulatory proteins or transcription factors (reviewed in ^10^). For instance, in mammalian cells, activated MYC associates with NUP153 and another NPC basket protein, TPR, at the nuclear periphery and triggers formation of a transcriptionally permissive environment that includes SAGA complex and MYC-target genes that mediate cell proliferation and migration ^93^. Second possible mechanism might be through establishment of a NUP153-mediated chromatin structure that provides optimal chromatin environment at putative CTCF and cohesin binding sites. CTCF-binding sites have characteristic features of chromatin structure including enrichment of histone modifications that include H3K4me3, H3K4me2, H3K4me1 and H2A.Z, and exhibit DNase I hypersensitivity ^88, 99, 100^. Thus, defining NUP153-interacting proteins and understanding their involvement in chromatin structure can provide valuable molecular insights on NUP153-mediated binding of chromatin architectural proteins.

Our findings are also relevant towards the understanding of diseases or cancer that underlie defects in chromatin-associated function of Nups. Several Nups, including NUP153 and NUP98 contain unstructured Phenylalanine-Glycine (FG)-repeats which mediate the phase separation during nucleocytoplasmic trafficking ^9, 101^. Structural chromosomal rearrangements or translocations of the FG-Nup genes result in formation of FG-Nup fusion proteins, which have been implicated in several hematologic malignancies including acute myeloid leukemia (AML) ^102^. For example, NUP98 fusion proteins are formed of a portion of the NUP98 FG repeats and a portion of a transcriptional regulator protein including CBP/P300, MLL, or DNA-binding proteins, such as HOX family members, HOXA9, HOXD13, and NSD1 ^103, 104, 105^ and influence expression of cancer-related genes. A recent report in *Drosophila* suggests that NUP98 forms a complex with several architectural proteins including CTCF ^22^. Thus, we propose that enhancer-specific regulation of chromatin structure and organization by mammalian NUP153 may apply to other FG-Nups and contribute to the gene regulatory mechanisms that underlie FG-Nup fusion protein associated cancers.

## METHODS

### Cell culture, plasmids, virus preparation and viral transduction

EL 16.7 female mouse ES cells (gift from J. Lee (Harvard)) and cell culture conditions have been described previously ^53^. ES cells were cultured on *γ*-irradiated mouse embryonic fibroblasts (MEFs) that were isolated from Tg(DR4)1Jae/J mice (The Jackson Laboratory). To transduce ES cells, control (scramble) or mouse NUP153 specific shRNA lentivirus particles (∼10^7^-10^8^ TU/ml) were added into 0.5 ml of complete ES cell medium containing dissociated ES cells (5×10^5^), LIF (ESGRO, Sigma-Aldrich) and Polybrene (4 μg/ml) (Sigma-Aldrich) and incubated overnight at 37°C. Next day, ES cells were dissociated and plated onto a 60-mm tissue culture dish (BD) containing *γ*-irradiated DR4 MEFs (1×10^6^), cultured for 24 hr in regular ES cell media followed by 2 days of selection using 2 μg/ml puromycin (Puro) (Sigma-Aldrich) and collected for subsequent analyses. HEK293T and HeLa cells were obtained from the American Tissue Collection Center (ATCC, Manassas, VA, USA) through the Duke University Cancer Center Facilities and were maintained in high glucose Dulbecco’s modified Eagle’s medium (DMEM) GlutaMAX supplemented with 10% FBS (Sigma-Aldrich), 1% penicillin/streptomycin, 1mM sodium pyruvate, 1% non-essential amino acids, and 3% HEPES. To generate FLAG-NUP153 overexpressing cells, HEK293T cells were transfected with Xfect reagent (Clonetech) with FLAG-mNUP153 or FLAG-hNUP153 cDNA vectors according to the manufacturer’s protocol. FLAG-hNUP153 (human) or FLAG-mNUP153 (mouse) expression vectors were constructed by amplifying full length human NUP153 or mouse NUP153 cDNA using human NUP153 cDNA (Origene, SC116943) or mouse NUP143 cDNA (ATCC, IMAGE clone ID: 6516328) clones, respectively. Amplified cDNA sequences were modified and cloned into *Bam*HI and *Xho*I sites of pCMV-3FLAG-6 vector (Agilent, 240200). To produce shRNA lentivirus particles, HEK293T cells were transfected with pMD2.G (Addgene #12259) and psPAX2 (Addgene #12260) vectors along with each shRNA lentiviral vector as previously described ^106, 107^. Viral supernatants were concentrated X100 using Lenti-X Concentrator (Clontech) according to the manufacturer’s protocol and stored at −80°C. All reagents were from Thermo Fisher Scientific, unless noted otherwise. All cells were cultured at 37°C with 5% CO_2_. Mouse husbandry and experiments were carried out as stipulated by the Duke University Institutional Animal Care and Use Committee (IACUC).

### Generation of NUP153 ES cell clones and NUP153 DamID-Seq

Mouse Nup153 cDNA (4.5 kb) (IMAGE clone ID: 6516328; ATCC) was modified and cloned in frame into *Kpn* I and *Xho* I sites in pIND-(V5)-EcoDam plasmid (gift from B. Van Steensel). To generate EcoDam or Nup153.EcoDam overexpressing ES cells, 10 μg of NUP153-(V5)-EcoDam-pIND or (V5)-EcoDam-pIND plasmid DNA were introduced into wild-type 16.7 female mouse ES cells ^53^ (1×10^7^) by electroporation (200 V, 1,050 µF) and stable clones were selected for 12 days using complete DMEM media supplemented with G418 (200 μg/ml) (Invitrogen). Positive clones were screened for EcoDam sequence by genomic DNA PCR using the primers: V5F, 5’-GGT AAG CCT ATC CCT AAC CCT C-3’; EcoDam_400R, 5’-AAC TCA CCG CGC AGA TTG TAA CG-3’ and by immunofluorescence as previously described ^108^. Mouse monoclonal *α*-V5 antibody was used in combination with rabbit polyclonal anti-IgG(H+L)-Alexa555 as a secondary antibody to detect V5-tagged Nup153.EcoDam fusion and EcoDam proteins (Figure 1SC). DamID was performed as described ^56^ with few modifications. Three 16.7 ES cell clones, expressing EcoDam (ED.B3) and NUP153.EcoDam fusion protein (NP.A2 and NP.D2) (E.Y., Y.S., R.I.S., Y.J., and J.T.L., manuscript in preparation) were used. Briefly, purified methyl PCR products were digested with *Dpn* II to remove adapter sequences from the fragment ends and 30 ng of PCR products were treated as DNA templates to prepare paired-end Solexa libraries as previously described ^109^. Genome Analyzer II (Illumina) was used to perform 2×36 cycles of paired-end sequencing. Sequencing reads from EcoDam overexpressing cells were used to normalize sequencing reads from NUP153.EcoDam overexpressing cells.

### Antibodies

Anti-CTCF (Millipore, 07-729), anti-SMC1A (Bethyl, A300-055A), anti-SMC3 (Abcam, ab9263), anti-FLAG (Sigma-Aldrich, F1804), anti-Nup153 (Abcam, ab24700), anti-GAPDH (Sigma-Aldrich, G9545), anti-Histone H3 (Abcam, ab1791), and anti-alpha-Tubulin (Santa Cruz, sc-5286) were used in western blot analysis. Anti-Rpb1 NTD (Cell Signaling, 14958), and anti-CTCF (Cell Signaling, 2899S) were used in ChIP analysis. Anti-Lamin B1 (Abcam, ab16048), anti-V5 tag (Thermo Fisher Scientific, R960-25), anti-IgG(H+L)-Alexa555 (Thermo Fisher Scientific, A-21427), and anti-IgG(H+L)-Alexa488 (Thermo Fisher Scientific, A-11008 and A-32723) were used in immunofluorescence.

### Immunoprecipitation (IP) assay

For IP assay, HEK293T cells that were transfected with FLAG-GFP or FLAG-NUP153 expression vector were lysed by sonication in IP lysis buffer (20 mM Tris-HCl, pH 7.9, 150 mM NaCl, 5 mM EDTA (pH:8.0), 1% Nonident P-40, 10% glycerol, 1 mM phenylmethylsulfonyl fluoride (PMSF), 1 mM DTT, protease inhibitor cocktail (Sigma-Aldrich)). After centrifugation, the supernatant was incubated with anti-FLAG M2 Affinity Gel beads (Sigma-Aldrich) at 4°C for 2 hr and the immune precipitates were subjected to western blotting. To prepare samples for the LC-MS/MS proteomics analysis, FLAG-NUP153 expression vector and mock transfected cells were lysed in elution buffer (10 mM PIPES, pH 6.8, 100 mM NaCl, 3 mM MgCl_2_, 0.3 M sucrose, 0.5% Triton X-100, 1 mM PMSF, 1 mM DTT, protease inhibitor cocktail (Sigma-Aldrich)) for 10 min on ice, the nuclear fraction containing pellet was collected by centrifugation 3 min, 500 x *g*, 4°C and was subjected to IP assay as described above. The immune precipitates were eluted by incubation with FLAG peptide (F4799) (Sigma-Aldrich) at room temperature for 15 min and were subjected to silver staining by using SilverXpress (Invitrogen) or utilized for LC-MS/MS proteomics analysis.

### LC-MS/MS Proteomics Analysis

Samples in 1X Laemmli Sample buffer (BIO-RAD, 1610737) were run on a NuPAGE 4-12% Bis-Tris Protein gel (Invitrogen, NP0336PK2) in NuPAGE MES SDS Running Buffer (Invitrogen, NP0002) for ∼5 min. The entire molecular weight range was excised and subjected to standardized in-gel trypsin digestion (http://www.genome.duke.edu/cores/proteomics/sample-preparation/documents/In gelDigestionProtocolrevised.pdf). Extracted peptides were lyophilized to dryness and resuspended in 12 uL of sample buffer (0.2% formic acid, 2% acetonitrile). Each sample was subjected to chromatographic separation on a nanoACQUITY UPLC (Waters) equipped with an ACQUITY UPLC BEH130 C_18_ 1.7 µm 75 µm I.D. X 250 mm column (Waters). The mobile phase consisted of (A) 0.1% formic acid in water and (B) 0.1% formic acid in acetonitrile. Following a 3 µL injection, peptides were trapped for 3 min on an ACQUITY UPLC M-Class Symmetry C_18_ Trap Column 5 µm 180 µm I.D. X 20 mm (Waters) at 5 µl/min in 99.9% A. The analytical column was then switched in-line and a linear elution gradient of 5% B to 40% B was performed over 30 min at 400 nL/min. The analytical column was connected to a SilicaTip emitter (New Objective) with a 10 µm tip orifice and coupled to a Q Exactive Plus mass spectrometer (Thermo Fisher Scientific) through an electrospray interface operating in a data-dependent mode of acquisition. The instrument was set to acquire a precursor MS scan from *m/z* 375-1600 at R=70,000 (target AGC 1e6, max IT 60 ms) with MS/MS spectra acquired for the ten most abundant precursor ions at R=17,500 (target ABC 5e4, max IT 60 ms). For all experiments, HCD energy settings were 27v and a 20 s dynamic exclusion was employed for previously fragmented precursor ions. Raw LC-MS/MS data files were processed in Proteome Discoverer (Thermo Fisher Scientific) and then submitted to independent Mascot search (Matrix Science) against a SwissProt database (*Human* taxonomy) containing both forward and reverse entries of each protein (20,322 forward entries). Search tolerances were 5 ppm for precursor ions and 0.02 Da for product ions using trypsin specificity with up to two missed cleavages. Carbamidomethylation (+57.0214 Da on C) was set as a fixed modification, whereas oxidation (+15.9949 Da on M) and deamidation (+0.98 Da on NQ) were considered dynamic mass modifications. All searched spectra were imported into Scaffold (v4.4, Proteome Software) and scoring thresholds were set to achieve a peptide false discovery rate of 1% using the PeptideProphet algorithm.

### Chromatin fractionation assay

The chromatin fractionation assay was performed as described previously ^48, 110, 111^ with minor modifications. Briefly, HEK293T cells (∼4 ×10^6^) were lysed in CSK buffer A (10 mM HEPES, pH 7.9, 10 mM KCl, 1.5 mM MgCl_2_, 340 mM sucrose, 0.1% Triton X-100, 10% glycerol, 1 mM PMSF, 1 mM DTT, protease inhibitor cocktail (Sigma-Aldrich, P8340)) for 10 min on ice. Total cell lysate (T) was separated into supernatant (S1, containing cytoplasmic proteins) and nuclei by centrifugation 3 min, 500 x *g*, 4°C. Nuclei were further lysed in CSK buffer B (3 mM EDTA, 0.2 mM EGTA, pH: 8.0, 1 mM DTT, 1 mM PMSF, protease inhibitor cocktail (Sigma-Aldrich, P8340)) and was incubated for 10 min on ice. The nuclear soluble fraction (S2, containing chromatin unbound proteins) and the nuclear insoluble fraction (P1) were separated by centrifugation 3 min, 500 x *g*, 4°C. The P1 fraction was resuspended in MNase buffer (10 mM Tris-HCl, pH 7.9, 10 mM KCl, 3 mM CaCl_2_, 300 mM sucrose, 1 mM PMSF, protease inhibitor cocktail (Sigma-Aldrich, P8340)) and chromatin was digested with 20 units of MNase (Thermo Fisher, EN0181) at room temperature for 15 min. The reaction was stopped by addition of EGTA (pH 8.0) at final concentration of 1 mM, followed by extraction with 250 mM ammonium sulfate at room temperature for 10 min. The chromatin-enriched fraction (S3, containing MNase-digested, chromatin associated proteins) and the nuclear insoluble fraction (P2, containing insoluble, nuclear membrane and nuclear matrix proteins) were collected from the supernatant and the pellet, respectively, by centrifugation 5 min, 1,700 x *g*, 4°C.

### Total RNA extraction, reverse transcription and real-time PCR

Total RNA was extracted from cells using TRIzol reagent (Invitrogen) according to the manufacturer’s protocol. For reverse transcription, cDNA was prepared using M-MLV Reverse Transcriptase (Thermo Fisher Scientific) with random hexamers (Sigma-Aldrich-Aldrich). Real-time PCR (qPCR) was performed using iTaq Universal SYBR Green Supermix (Bio-Rad) with specific primer sets indicated in Table S1. Relative gene expression was calculated by the relative standard curve method. *GAPDH* expression was used to normalize data.

### IEG transcription induction in HeLa cells

HeLa cells (1×10^6^) were transduced with the control (scramble) or human NUP153-specific shRNA lentivirus particles overnight at 37°C followed by selection in medium containing Puromycin (Puro, 2 μg/ml) (Sigma-Aldrich) for 48 hr. To collect cells at the basal (minus EGF) IEG state, cells were pre-cultured in DMEM supplemented with 0.1% FBS (Sigma-Aldrich) for 24 hr, followed by EGF (50 ng/ml) (Sigma-Aldrich, E9644) treatment for 15, 30, 60, 90 and 120 min. For the recue experiments, HeLa cells were transfected with control (scramble) or NUP153-specific shRNA vectors along with FLAG-hNUP153 expression vector using Xfect transfection reagent (Clontech) according to the manufacturer’s protocol. At the 16 hr time point, culture medium was replaced with Puro (2 μg/ml) containing medium and cells were incubated in this medium for 24 hr, followed by incubation in Puro-free medium for another 24 hr. To induce IEG transcription, cells were subjected to EGF treatment as described above.

### Chromatin immunoprecipitation (ChIP) assay

ChIP experiments were performed as previously described ^112^. Briefly, mouse ES cells (∼2×10^6^) were cross-linked with 1% formaldehyde at room temperature for 10 min and the reaction was stopped by adding glycine (final concentration, 125 mM). Cross-linked cells were treated with a hypotonic buffer (10 mM HEPES-NaOH, pH 7.9, 1.5 mM MgCl_2_, 10 mM KCl, 0.2% NP-40, 1 mM DTT, 1 mM PMSF, protease inhibitor cocktail (Sigma-Aldrich) at 4°C for 10 min, and collected in a sample tube. The cells were lysed in ChIP lysis buffer (50 mM Tris-HCl, pH 7.9, 10 mM EDTA, 1% SDS, 1 mM PMSF, protease inhibitor cocktail (Sigma-Aldrich) on ice for 10 min and were subjected to sonication to shear the chromatin to 200-1,000 bp-long DNA fragments, and were diluted with 9x volumes of ChIP dilution buffer (16.7 mM Tris-HCl, pH 7.9, 167 mM NaCl, 1.2 mM EDTA, 1.1% Triton X-100, 1 mM PMSF). The lysate was incubated with each antibody at 4°C overnight with rotation, and DNA-protein complexes were pulled down using Protein A/G agarose beads (Thermo Fisher, 20423). Agarose beads were washed with the following buffers: low-salt wash buffer (20 mM Tris-HCl, pH 7.9, 100 mM NaCl, 2 mM EDTA, pH:8.0, 0.1% SDS, 1% Triton X-100), high-salt wash buffer (20 mM Tris-HCl, pH 7.9, 500 mM NaCl, 2 mM EDTA, 0.1% SDS, 1% Triton X-100), and the LiCl buffer (10 mM Tris-HCl, pH 7.9, 250 mM LiCl, 1% NP-40, 1% deoxycholate acid, 1 mM EDTA, pH 8.0), and twice with TE buffer. DNA-protein complexes were eluted by incubating the beads in the elution buffer (1% SDS, 0.1 M NaHCO_3_) for 15 min at room temperature with rotation. Samples were reverse cross-linked by overnight incubation at 65°C in 200 mM NaCl followed by proteinase K (20 μg/ml) treatment at 55°C for 2 hr. DNA was purified using MinElute PCR purification kit (QIAGEN). DNA samples were used to measure occupancy of each protein across genes using gene-specific primer sets (Table S1) by real-time PCR as described above.

### Immunostaining and DNA FISH

For sequential Lamin B1 immunostaining and c-FOS DNA FISH, HeLa cells (5.5×10^3^) were grown on 12-well glass slides (Invitrogen) overnight at 37°C, and IEG transcription was induced as described above. Immunostaining was performed as previously described ^108^. Briefly, fixed cells were subjected to immunostaining using anti-Lamin B1 (Abcam, ab16048) antibody (1:400) at 4°C overnight, washed three times in wash buffer (1xPBS/0.2% Tween-20 buffer) at room temperature for 5 min each, and incubated with goat polyclonal anti-IgG(H+L)-Alexa 488 secondary antibody (1:500) for 1 hr at room temperature. To remove excess secondary antibody, cells were washed three times in wash buffer for 5 min each. Slides were mounted using Vectashield mounting medium containing DAPI (Vector Labs). Slides were imaged using a Leica DM5500B microscope, and a Leica DFC365 FX CCD camera, image positions were recorded and slides were washed in 1xPBS/0.2% Tween 20 to remove the mounting medium and re-fixed in 4% paraformaldehyde (Electron Microscopy Sciences) prior to DNA FISH experiment. To detect DNA signal at the *c-FOS* locus by FISH, BAC clone (RP11-293M10) (CHORI) was fluorescently labeled using Cy3-dUTP (ENZO) and nick translation kit according to the manufacturer’s instructions (Sigma-Aldrich). Human Cot-1 (DNA) (Thermo Fisher Scientific) (10 μg per 2 μg of nick translated BAC vector) was included to block background DNA signal, the probe was precipitated by NaOAc-EtOH precipitation and the probe was re-suspended in 50 μl of hybridization buffer (50% Formamide, 2xSSC, 2 mg/ml BSA (Sigma-Aldrich), 10% Dextran sulfate-500K (Sigma-Aldrich)) generating ∼40 ng/μl labeled DNA probe. DNA FISH was performed as previously described ^108^. Hybridization was performed using ∼200 ng DNA probe at 37°C overnight in a humidified chamber. DNA FISH images at the recorded positions were obtained with a Leica DM5500B microscope, a Leica DFC365 FX CCD camera, and analyzed using Fiji software. Distribution of *c-FOS* locus distance to nuclear periphery was measured in control and NUP153 KD HeLa cells at the indicated time points. Cumulative frequencies at a normalized distance (ND) of 0.0-0.12 are shown (Figure 7). Frequency of *c-FOS* distribution at ND 0.0-0.45 are shown in Figure S7. ND= (*c-FOS* locus to periphery distance)/(cell diameter (d)), where d=(2xnuclear area/*π*)^0.5^. \**p*<0.05; \*\*\**p*<0.001; the Kolmogorov–Smirnov (KS)-test was applied to calculate significance. To determine the cellular distribution of FLAG-NUP153 in HEK293T cells, FLAG-NUP153 transfected HEK293T cells (5.5×10^3^) were cultured on glass coverslips, and immunostaining was performed using anti-FLAG M2 (Sigma-Aldrich, F1804) antibody (1:250) as described above.

### Poly(A)^+^ RNA FISH and alkaline phosphatase staining

Poly(A)^+^ RNA FISH was performed by using 5’ Cy3-labled oligo-dT 50mer (Sigma-Aldrich) as previously described ^66^. Briefly, hybridization was performed using 0.5 μg oligo-dT probe per sample overnight at 37°C in a humidified chamber. Following hybridization, cells were washed twice for 15 min at 42°C with 2XSSC, and once for 15 min at 42°C in 0.5XSSC. Slides were mounted using mounting medium containing DAPI (Vector labs) and cells were imaged by fluorescence microscopy. Alkaline phosphatase staining was performed using Red Alkaline Phosphatase Substrate kit (Vector labs, SK-5100) according to the manufacturer’s protocol. Bright field images were taken using Leica EC3 color camera attached to Leica DM5500B microscope.

### Nuclear transport assay

Hela cells were co-transfected with Rev-Glucocorticoid Receptor-GFP (RGG) expression vector (Gift from K. Ullman (University of Utah)) ^113^ and control (scrambled) or hNUP153-specific shRNA vectors using Xfect reagent. Import and export assays were performed as previously described ^81^. Briefly, for import assay, transfected HeLa cells were grown overnight on 12-well glass slides at 37°C and treated with 250 nM dexamethasone (Dex) (Sigma-Aldrich, D4902) to induce RGG nuclear import for the indicated times. For the export assay, 120 min Dex-treated cells were washed with 1xPBS (pH:7.2) and cultured in fresh culture medium for the indicated times. At the end of each time point, cells were fixed using 4% paraformaldehyde and mounted using DAPI containing mounting medium (Vector labs) Images were obtained with a Leica DM5500B microscope, a Leica DFC365 FX CCD camera, and examined to calculate percentage of cells with nuclear GFP-RGG signal.

### RNA-Seq

Total RNA quality and concentration was assessed on a 2100 Bioanalyzer (Agilent Technologies) and Qubit 2.0 (ThermoFisher Scientific), respectively. Total RNA (RIN value ≥ 8) from control and two NUP153 KD ES cells were depleted of ribosomal RNA using the Illumina Ribo-zero Gold kit and converted into RNA-seq libraries using the Illumina Total RNA-seq kit. Libraries were indexed using a dual indexing approach allowing for multiple libraries to be pooled and sequenced on the same sequencing flow cell of an Illumina HiSeq 4000 sequencing platform. Before pooling and sequencing, fragment length distribution and library quality was first assessed on a Fragment Analyzer (Agilent). All libraries were eventually pooled in equimolar ratio and sequenced. Libraries were sequenced at 50bp single-end on the Illumina HiSeq 4000 instrument. About 110×10^6^ reads per sample were generated. Once generated, sequence data was demultiplexed and Fastq files generated using Illumina’s Bcl2Fastq v2 conversion software.

### ChIP-Seq

ChIP DNA samples were quantified using the fluorometric quantitation Qubit 2.0 system (ThermoFisher Scientific). ChIP-Seq libraries were prepared using the Roche Kapa BioSystem HyperPrep Library Kit to generate Illumina-compatible libraries. During adapter ligation, dual unique indexes were added to each sample. Resulting libraries were cleaned using SPRI beads and quantified using Qubit 2.0. Fragment length distribution of the final libraries was assessed on a Fragment Analyzer (Agilent). Libraries were then pooled into equimolar concentration and sequenced on an Illumina HiSeq 4000 instrument. Sequencing was done at 50bp single-end and generated about 110×10^6^ reads per sample. Sequence data was demultiplexed and Fastq files generated using Illumina’s Bcl2Fastq v2 conversion software.

### Analyses of RNA-Seq, ChIP-Seq data and TAD boundaries

#### RNA-Seq data analyses

RNA-Seq reads were trimmed by Trim Galore (0.4.1, with -q 15) and then mapped with TopHat (v 2.1.1, with parameters --b2-very-sensitive --no-coverage-search and supplying the UCSC mm10 known gene annotation). The ERCC spike-in sequences were mapped separately. Gene-level read counts were obtained using the featureCounts (v1.6.1) by the reads with MAPQ greater than 30. Bioconductor package RUVseq (v 1.16.0) was used to normalize the read counts and edgeR (v 3.24.0) was employed for differential expression analysis. Fold change greater than 1.5 and false discovery rate (FDR) less than 0.05 was used to filter the significant differentially expressed genes.

### ChIP-Seq

#### Definition of regulatory regions

Several analyses in the manuscript rely on ChIP-Seq analyses across different regulatory regions namely enhancers and promoters. Below we describe how these regulatory regions were defined.

#### Promoters

Promoters are defined by gene start sites downloaded from UCSC Genome Browser goldenPath/mm10/database/knownGene. Active promoters were defined by the Fragments Per Kilobase of transcript per Million mapped reads (FPKM), which is calculated by cufflinks(v 2.1.1), greater than 1 in control RNA-seq. Inactive promoters were defined by FPKM no greater than 1. Chromatin structure at the transcriptionally active vs inactive TSS was validated using previously published H3K4me3 and H3K27me3 ChIP-Seq, respectively (GEO: GSE36905) ^57^.

#### Enhancers

Enhancers were defined by utilizing the previously published ChIP-Seq data sets and determining the overlapping region of peaks with at least two enhancer specific markers including CBP/P300 (GEO: GSE29184), H3K4me1 (GEO: GSE25409) or H3K27Ac (GEO: GSE42152).

#### TAD boundaries

TAD boundaries were defined by utilizing the previously published Hi-C data and TAD boundary coordinates reported ^61^.

#### Determining NUP153-CTCF or NUP153-cohesin co-occupied sites

The overlap between NUP153 DamID peaks and CTCF or SMC3 ChIP peaks were defined using control (scramble shRNA) samples and were called by utilizing the Bioconductor package ChIPpeakAnno (v. 3.19.5) with a maximal gap of 5kb. The overlapping sites are referred to as co-occupied sites.

### ChIP-Seq data analyses

ChIP-Seq reads were trimmed by Trim Galore (0.4.1, with -q 15) and then mapped with bowtie2 (2.2.5, with parameters --very-sensitive) to mouse genome (UCSC mm10). The mapped reads were filtered by MAPQ greater than 30 by samtools (v 1.5) and duplicated reads were removed by picard (v 1.91). The peaks were called by MACS2 (v 2.1.0, with - -pvalue 1e-5). The read coverages were quantified by the signal in reads per million per base pair https://github.com/BradnerLab/pipeline/blob/master/bamToGFF.py with parameters -m 500 -r -d. Metagene plots were used to display the average ChIP-seq signal across related regions of interest for enhancers and TSS separately. The average profile (metagene) was calculated by the mean of ChIP-seq signal profiles across the related regions of interest. For each metagene plot, the profile is displayed in rpm/bp in a ±2.5 or 5 kb region centered on the regions of interest. The number of enhancers or TSS were noted in the title of plots.

### DamID-Seq

DamID-Seq reads were mapped with bowtie2 (2.2.5, with parameters --very-sensitive) to mouse genome (UCSC mm10). The mapped reads were filtered by MAPQ greater than 30 by samtools (v 1.5) and filtered by GATC in 5’ ends. The peaks were called by MACS2 (v 2.1.0, with -q 0.05). To determine distribution of NUP153-DamID peaks across the genetic elements in mouse ES cells we used the following criterion. Promoters (-2kb from TSS to +100 bp from TSS); GB (+100bp from TSS to +1kb from TTS); Intergenic sites (< −2kb from TSS and >+1kb from TTS). TSS, transcription start site; GB, gene body; TTS, transcription termination site.

### Hi-C analyses

Mouse ES cell normalized 40kb HiC Matrices (mm9) were downloaded from http://chromosome.sdsc.edu/mouse/hi-c/download.html. The Hi-C 2D map were plotted by R/Bioconductor package trackViewer (v. 1.23.2)

### Examining IEG chromatin structure in HeLa cells

ENCODE HeLa-S3 ChIP-Seq data sets ^82^ for POL II (GEO: GSM733759), CTCF (GEO: GSM733785), RAD21 (GEO: GSM935571), CBP/P300 (GEO: GSM935553), H3K4me1 (GEO: GSM798322), H3K27Ac (GEO: GSM733684), and H3K4me3 (GEO: GSM733682) were utilized to examine chromatin structure across the JUN and EGR1 genes (Figure S6B) using Human hg19 as a reference genome.

### Statistical analysis

Quantitation of data was performed using the following statistical tests. Significance of the difference between control and knockdown cells for variables was analyzed with parametric Student’s t-test. The nonparametric Kolmogorov–Smirnov (KS)-test was applied to calculate significance between the control and knockdown cells during the analyses of *c-FOS* locus spatial positioning with respect to nuclear periphery in a time course dependent manner.

### Reporting summary

Further information on research design is available in the Nature Research Reporting Summary linked to this article.

### Data availability

Gene expression profiles, DamID-Seq and ChIP-Seq datasets have been deposited at GEO with the accession code GSE135647. Proteomics data have been deposited to the ProteomeXchange Consortium via the PRIDE partner (https://www.ebi.ac.uk/pride/) repository with the Project ID: PXD015441. The source data underlying Figs. 1a-b, 1d, 4a-d, 5, 6a-d, 7a-b and Supplementary Figs. 1a, 5a-b, 7 are provided as a Source Data file. All other relevant data supporting the key findings of this study are available within the article and its Supplementary Information files from the corresponding author upon request.

### Code availability

The custom analysis pipelines for all genomic analyses are available upon request with no restrictions.

## ACKNOWLEDGEMENTS

We are grateful to the members of the Yildirim lab for critical discussions and feedback. We thank L. Birnbaumer, and S. Namekawa for critical reading and feedback on the manuscript. We are grateful to J.T. Lee at whose lab the NUP153-DamID constructs were generated by E. Yildirim and DamID-seq data was produced. We acknowledge B. Van Steensel for the DamID plasmid, K. Ullman for the REV-GFP-GR plasmid, C. Gersbach for the pMD2.G and psPAX2 plasmids, J. Black for assistance in virus preparation. We thank E. Soderblom at the Duke Proteomics Shared Resource for assistance in the Mass-spec experiment, N. Devos at the Duke Genome Sequencing Shared Resource for assistance in ChIP-, RNA-Seq library preparation and sequencing, B. Cantor at the Duke Viral Shared Resource for assistance in virus preparation, and Duke Computing Cluster for computing support. This work was supported by the Whitehead Scholar Award (E.Y.).

## AUTHOR CONTRIBUTIONS

E.Y. conceptualized and supervised the study. S.K., and E.Y. designed the experiments. S.K. performed all aspects of experiments associated with identification of NUP153-interacting proteins, the role of NUP153 in regulation of IEG transcription and chromatin structure in HeLa cells. Y.S. carried all aspects of mouse ES cell ChIP assays and Poly (A)+ RNA-FISH in collaboration with S.K. and E.Y. J.S. conducted and analyzed GR import/export assays in collaboration with S.K. and E.Y. J.O. analyzed RNA-, DamID-, Hi-C, and ChIP-Sequencing data together with S.K. and E.Y. E.Y. performed Lamin B1 immunostaining and c-FOS DNA-FISH and imaging. S.K. performed FLAG-NUP153 immunostaining and imaging. S.K., J.O., Y.S., J.S., and E.Y. analyzed the experiments. S.K. and E.Y. wrote the paper with input from all authors.

## MATERIALS & CORRESPONDENCE

Further information and requests for resources and reagents should be directed to and will be fulfilled by the Lead Contact, Eda Yildirim.

## SUPPLEMENTAL FIGURE LEGENDS

**Figure S1:**
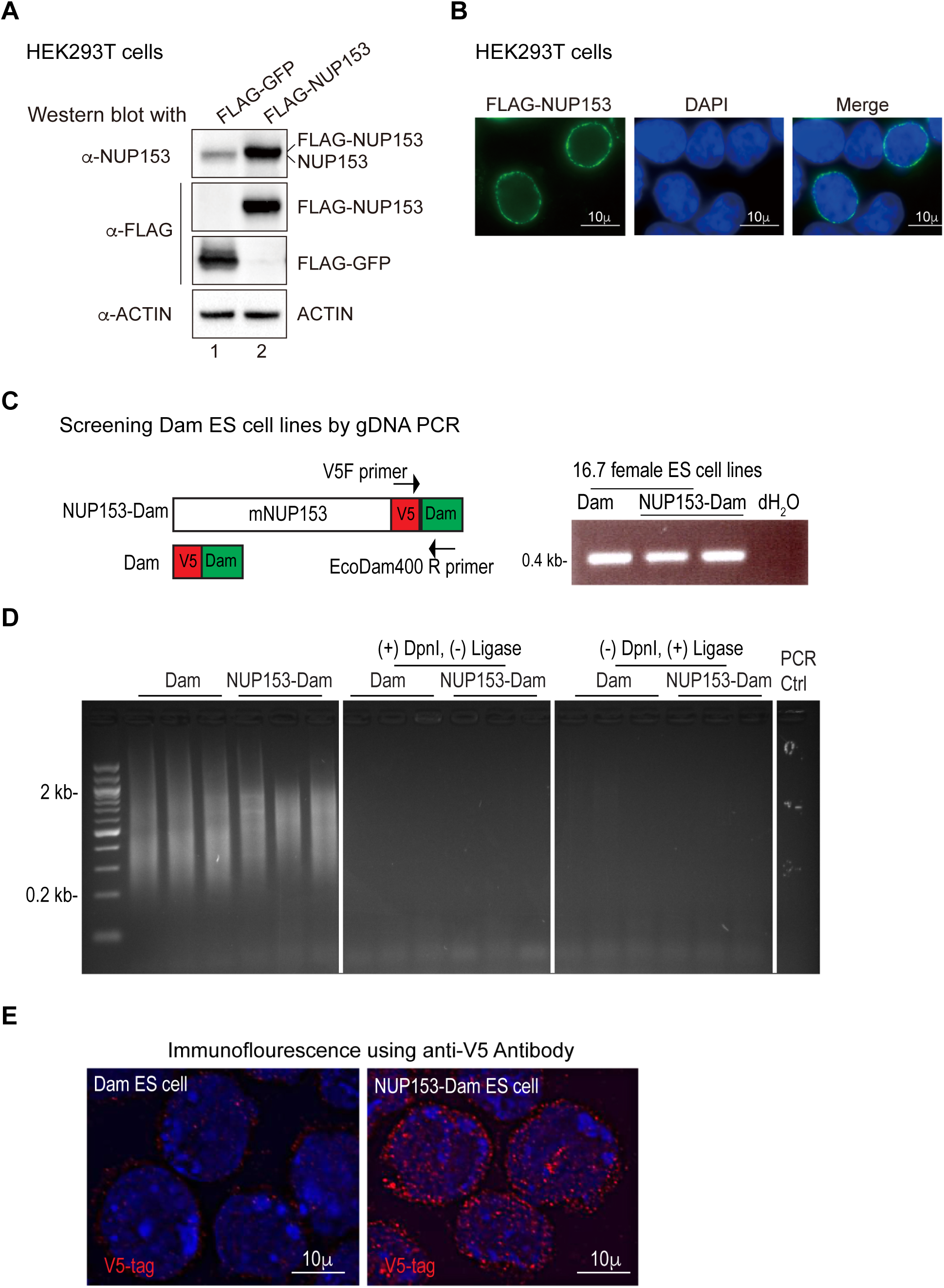
FLAG-NUP153 expression in HEK293T cells and generation of NUP153-Dam mouse ES cell lines. **Related to** Figure 1, 2, 3. (A) Western blot showing validation of FLAG-NUP153 expression. Whole cell extracts were prepared from FLAG-GFP- or FLAG-NUP153-expressing HEK293T cells and were subjected to western blot using anti-NUP153, and anti-FLAG antibodies as indicated. ACTIN was used as an internal control. (B) Cellular localization of FLAG-NUP153 protein in HEK293T cells was determined by immunostaining. Scale, 10μm. (C) Mouse NUP153-cDNA (4.5 kb) (ATCC) was cloned into *Kpn* I and *Xho* I sites in pIND-(V5)-EcoDam plasmid ^56^ to generate NUP153-Dam plasmid. Female mouse ES cell line 16.7 were electroporated using NUP153-Dam or Dam only (pIND-(V5)-EcoDam) plasmids. ES clones were screened by genomic DNA (gDNA) PCR using a primer pair that amplifies a 0.4 kb fragment across V5-tag and Dam sequences. (D) Several ES cell clones were screened for Dam activity by gDNA PCR as previously described ^56^. (E) Expression of NUP153-Dam fusion protein was determined by performing immunofluorescence using anti-V5 antibody in Dam only and NUP153-Dam mouse ES cells. Scale, 10μm.

**Figure S2:**
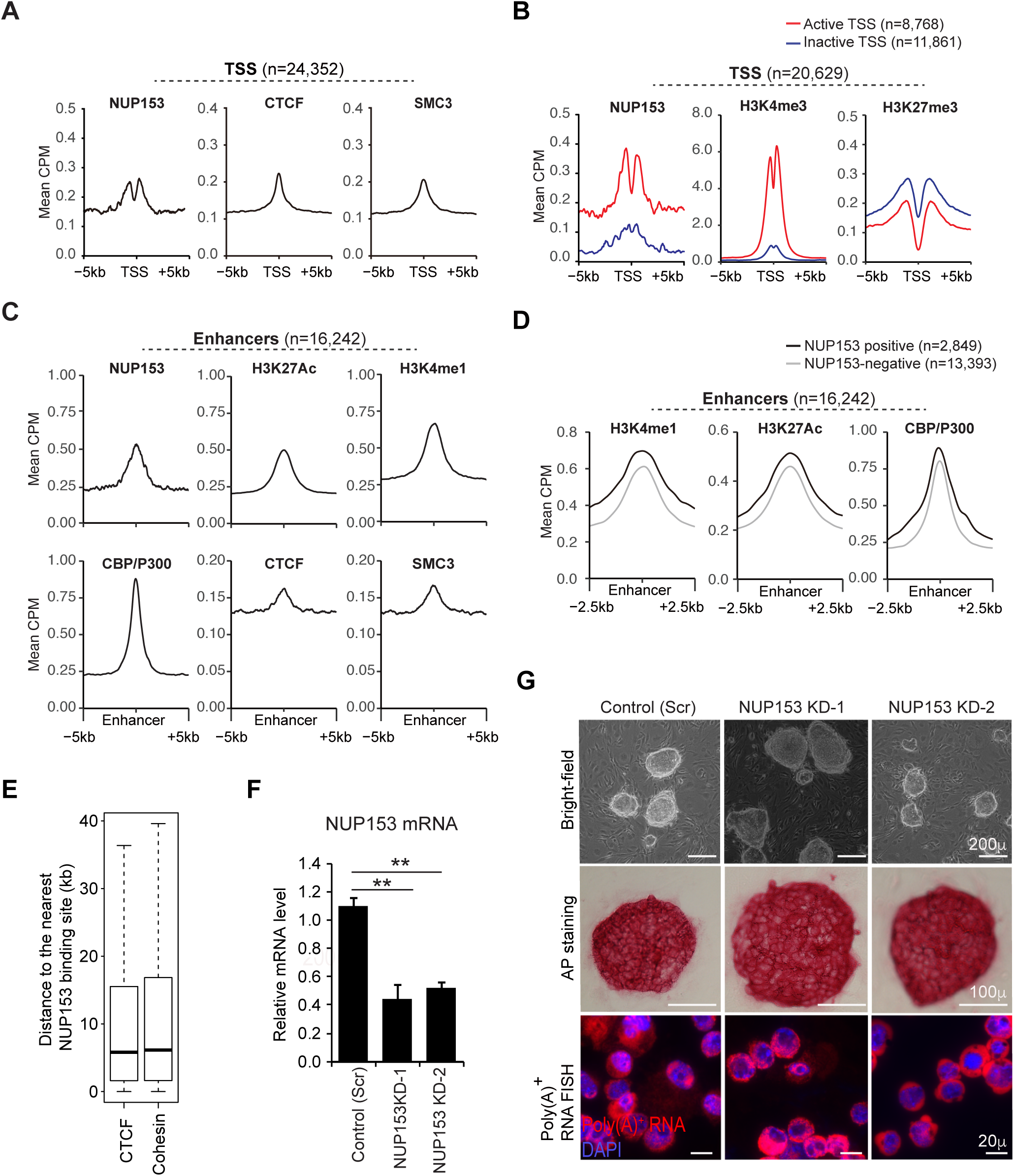
Distribution of NUP153, CTCF, SMC3 and histone modifications across genetic elements in control (WT) mouse ES cells, and characteristics of NUP153 deficient mouse ES cells. **Related to** Figure 2 **and** 3. (A) Metagene profiles showing mean CTCF, SMC3 and NUP153 binding at TSS (+/-5kb) (n=24,352). (B) Metagene profiles showing mean CTCF, SMC3 and NUP153 binding +/− 5kb of transcriptionally active (n=8,768) and inactive (n11,861) TSS. (C) Metagene profiles showing mean NUP153, H3K27Ac ^59^, H3K4me1 ^58^, CBP/P300 ^60^, CTCF, and SMC3 binding at enhancers (n=16,242)(+/-5kb). (D) Metagene profiles showing distribution of H3K27Ac ^59^, H3K4me1 ^58^, CBP/P300^60^ at NUP153-positive (n=2,849) and NUP153-negative (n=13,393) enhancers (+/− 2.5kb) in control ES cells. (E) Distribution of CTCF or cohesin binding sites was evaluated to determine the median of their distance to the nearest NUP153 binding sites in mouse ES cells. CTCF and cohesin binding sites exhibit a median of ∼5 kb distance to the nearest NUP153 binding sites. (F) Real time RT-PCR showing relative NUP153 mRNA levels in control (scramble) and NUP153 shRNA (KD-1, KD-2) virus transduced ES cells. *GAPDH* mRNA level was used to normalize mRNA levels. Student’s t-test was applied to calculate significance. Mean mRNA levels ± standard error mean. \*\**p*<0.01. (G) Top panel, bright-field microscopy images showing typical pluripotent ES cell morphology of NUP153 deficient and control ES cells. Scale 200μm; middle panel, immunostaining for alkaline phosphatase (AP) activity validates pluripotency of control and NUP153 KD ES cells. Scale 100μm; bottom panel, oligo(dT)50-Cy3 RNA FISH for Poly(A)^+^ RNA was performed in control and NUP153 KD ES cells to assess Poly(A)^+^ RNA export. No significant defect in Poly(A)^+^ RNA export was detected in NUP153 deficient cells. Scale 200μm. Experiments repeated at least three times.

**Figure S3:**
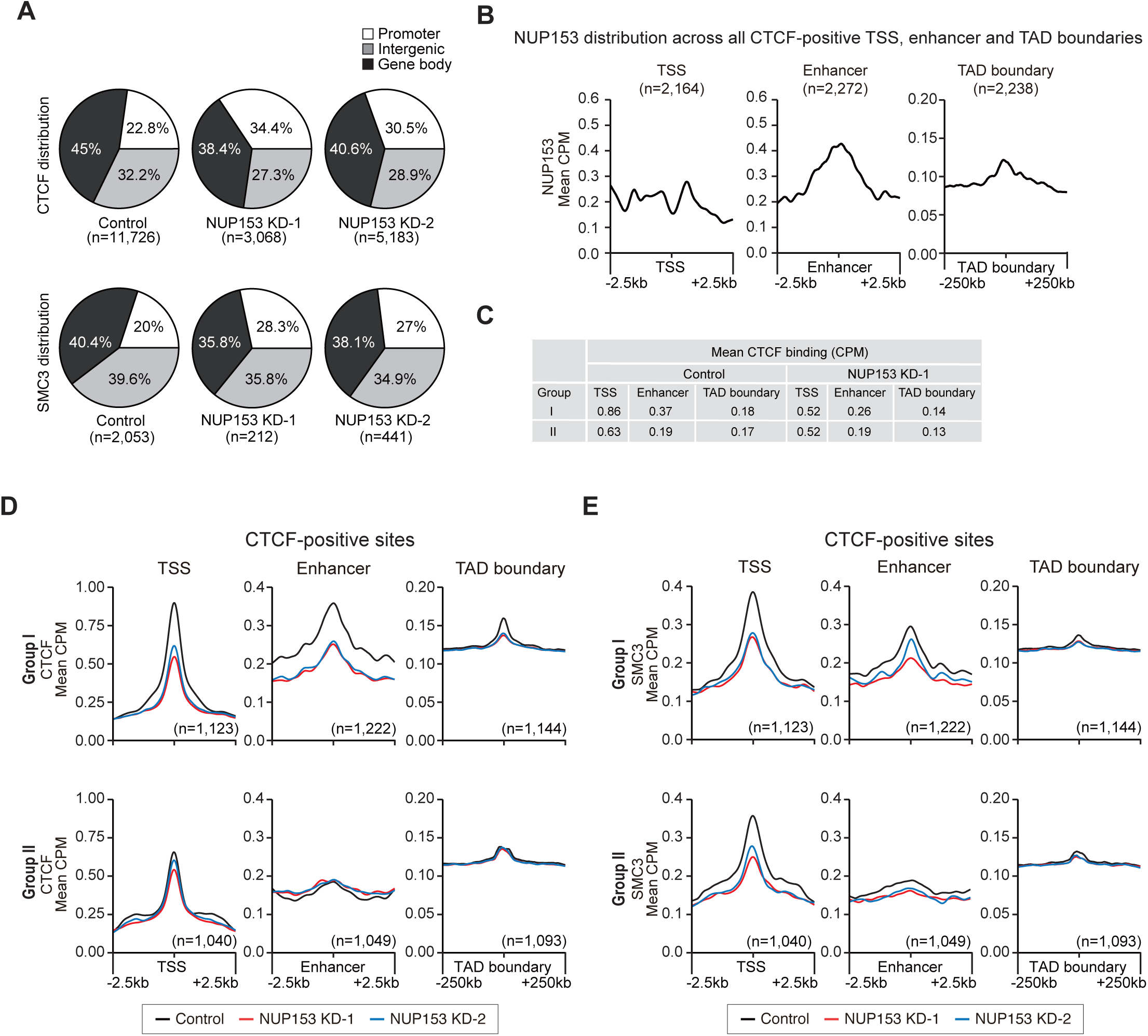
Distribution of CTCF, SMC3 and NUP153 across genetic elements and CTCF-positive sites in mouse ES cells. **Related to** Figure 2 **and** 3. (A) Distribution of CTCF and SMC3 sites across the indicated genetic elements in control and NUP153 KD-1 and KD-2 mouse ES cells. (B) Metagene profiles showing mean NUP153 binding at CTCF-positive TSS, enhancer and TAD boundaries. (C) Table showing mean CTCF binding at CTCF-positive TSS, enhancer and TAD boundaries in control and NUP153 KD ES cells. (D-E) Metagene profiles showing mean CTCF (D) and SMC3 (E) binding in control versus NUP153 KD cells at CTCF-positive Group I and Group II TSS, enhancer and TAD boundaries. Number of CTCF-positive sites for each Group is as indicated.

**Figure S4:**
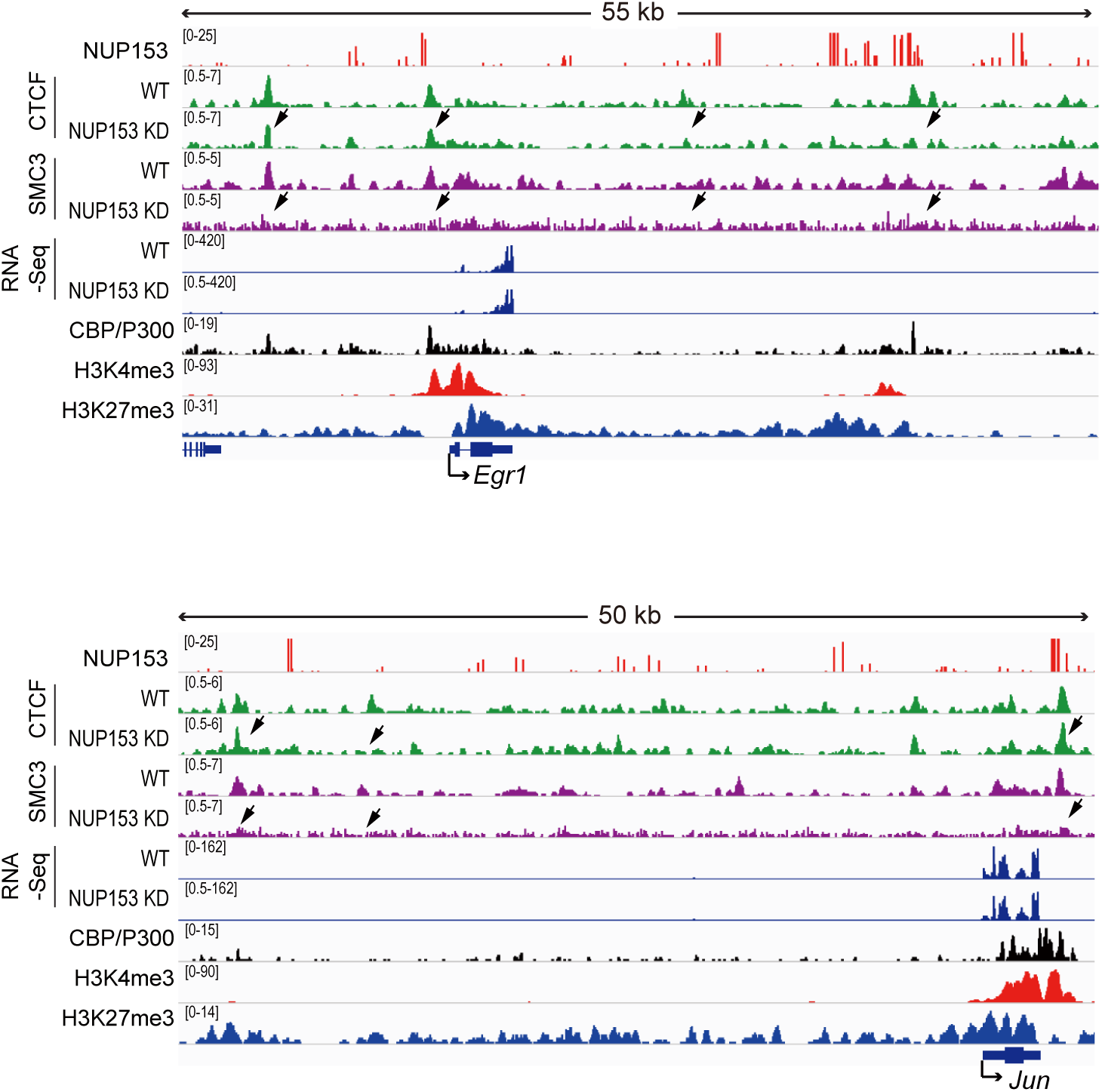
IEG loci are NUP153 targets in mouse ES cells. **Related to** Figure 3. CBP/P300, H3K4me3, H3K27me3, NUP153, CTCF, SMC3 ChIP-Seq, NUP153 DamID-Seq and RNA-Seq tracks are shown for IEG loci, *Egr1* (top panel), and *Jun* (bottom panel) in control and NUP153 KD ES cells. CBP/P300 ^60^, H3K4me3, H3K27me3 ChIP-Seq data ^57^ were previously published

**Figure S5:**
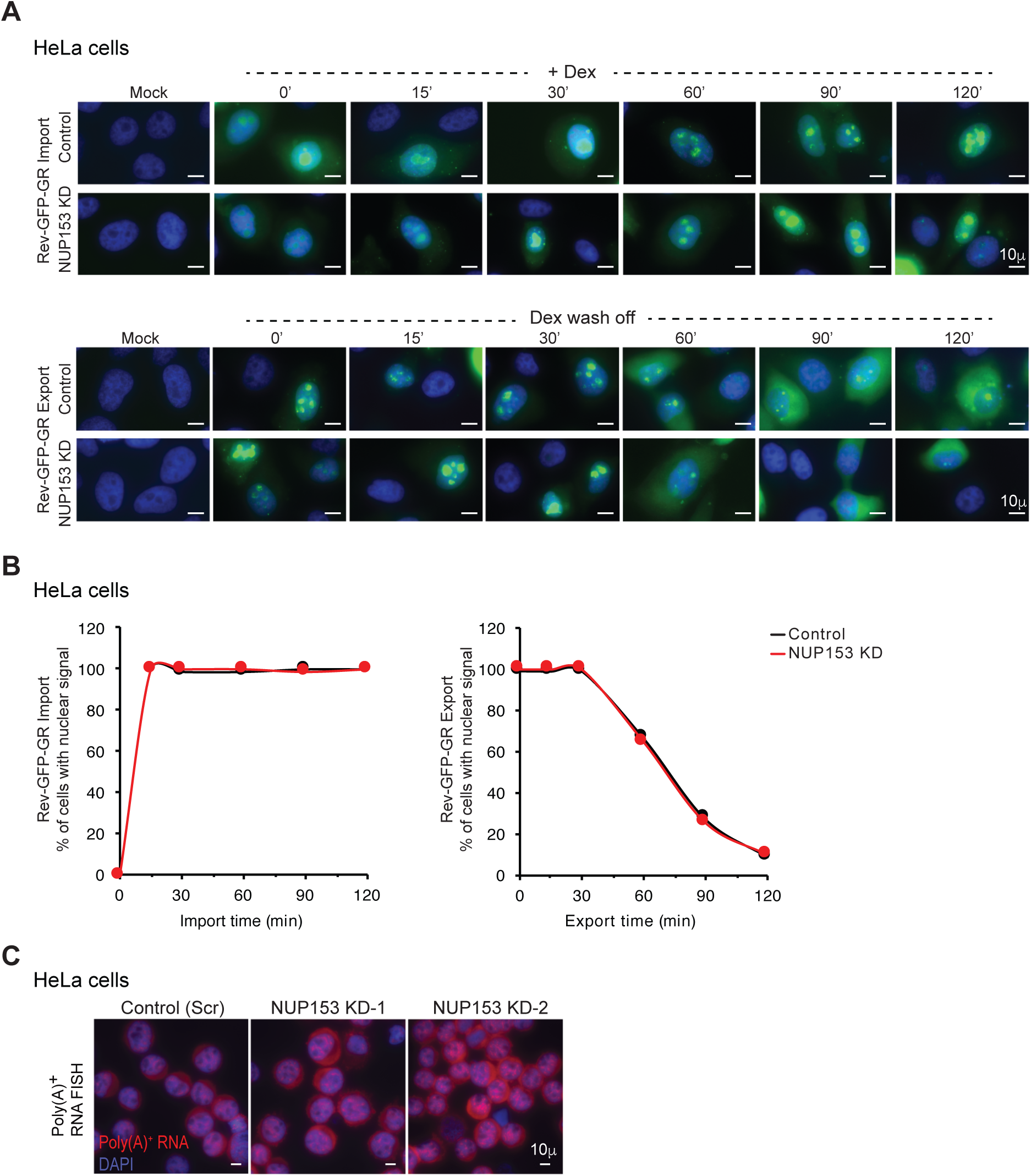
Analyses of NUP153 deficient HeLa cells for nucleocytoplasmic trafficking. **Related to** Figure 4. (A) Nuclear import and export were tracked using the dexamethasone responsive REV-GFP-GR construct (see Methods for more detailed information). Shown are representative images of control and NUP153 KD HeLa cells after dexamethasone (Dex) treatment or wash off at the indicated time points. To evaluate GR import, cells were treated with 250 mM Dex at the indicated times (n=68-84). To evaluate GR export, Dex was washed off after 120 mins (considered zero (0) time point) (n=75). (B) Graphs showing nuclear import of REV-GFP-GR after Dex treatment (left) and export of REV-GFP-GR after Dex wash off (right). Values were calculated based on % of cells which show nuclear GR-GFP signal after Dex treatment, or Dex wash off at the indicated time points. Scale bar, 10μm. (C) Oligo(dT)50-Cy3 RNA FISH in control and NUP153 KD HeLa cells was performed to evaluate Poly(A)^+^ RNA export. Scale 10μm.

**Figure S6:**
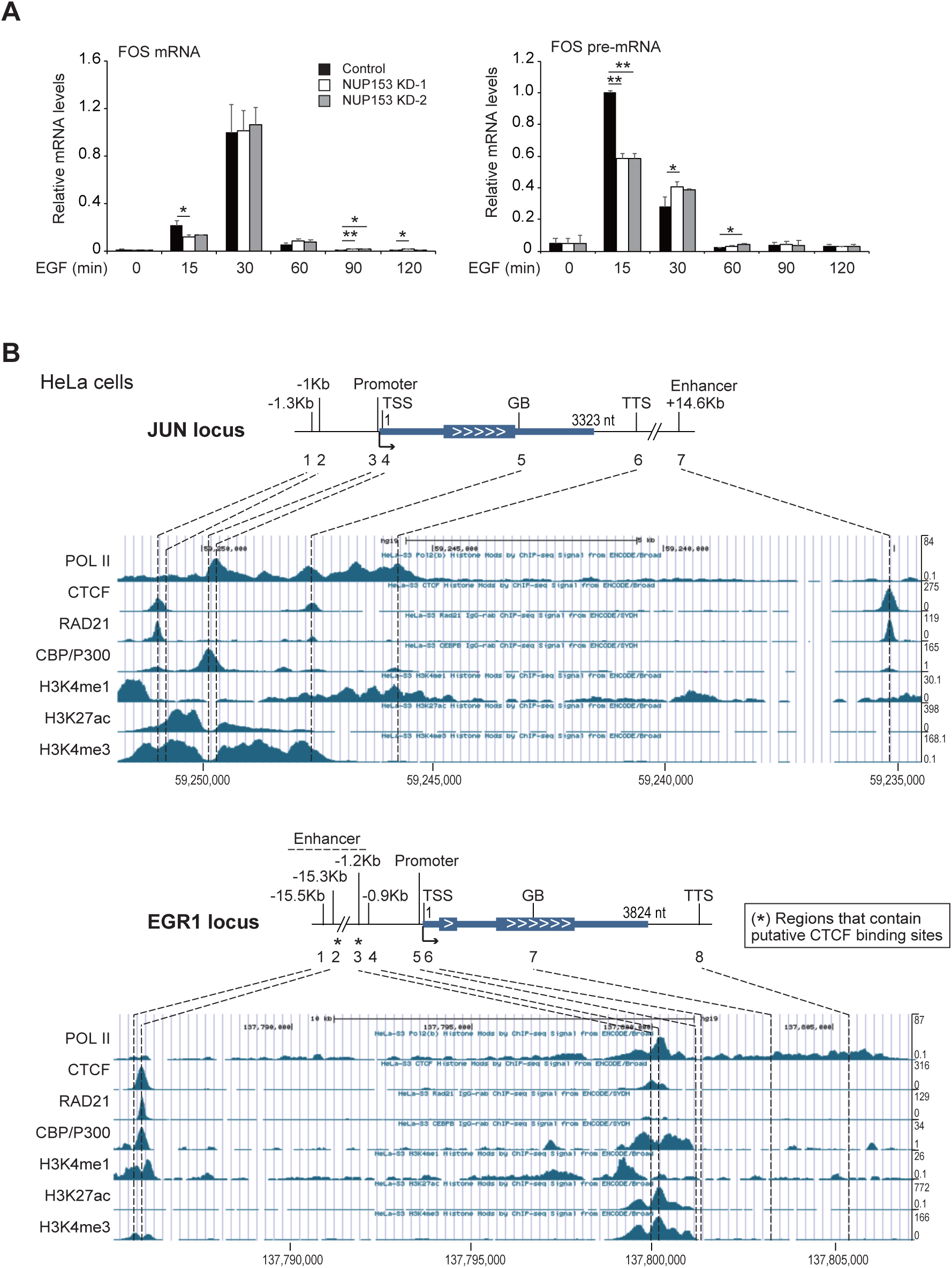
Transcription and chromatin structure at IEG loci in HeLa cells. **Related to** Figure 4 and 5. (A) Real-time RT-PCR showing relative mRNA and nascent mRNA levels for *c-FOS* gene in control and NUP153 KD HeLa cells in a time course dependent manner. *GAPDH* was used to normalize mRNA levels. Values are mean ± standard deviation. Student’s t-test was applied to calculate significance. \**p*<0.05; \*\**p*<0.01. Experiments were repeated at least 3 times. (B) UCSC browser snapshots showing distribution of POL II, CTCF, cohesin subunit, RAD21, CBP/P300, and histone modifications, H3K4me1 and H3K27Ac, H3K4me3, across the *JUN* (top panel) and *EGR1* (bottom panel) loci in HeLa-S3 cells based on ENCODE ChIP-Seq datasets (see Methods for details on GEO information for each data set). ChIP-Seq read numbers are indicated at the right *y*-axis per data set. Human hg19 reference genome was used to analyze data sets. Asterisk (*) denotes sites that contain putative CTCF binding.

**Figure S7:**
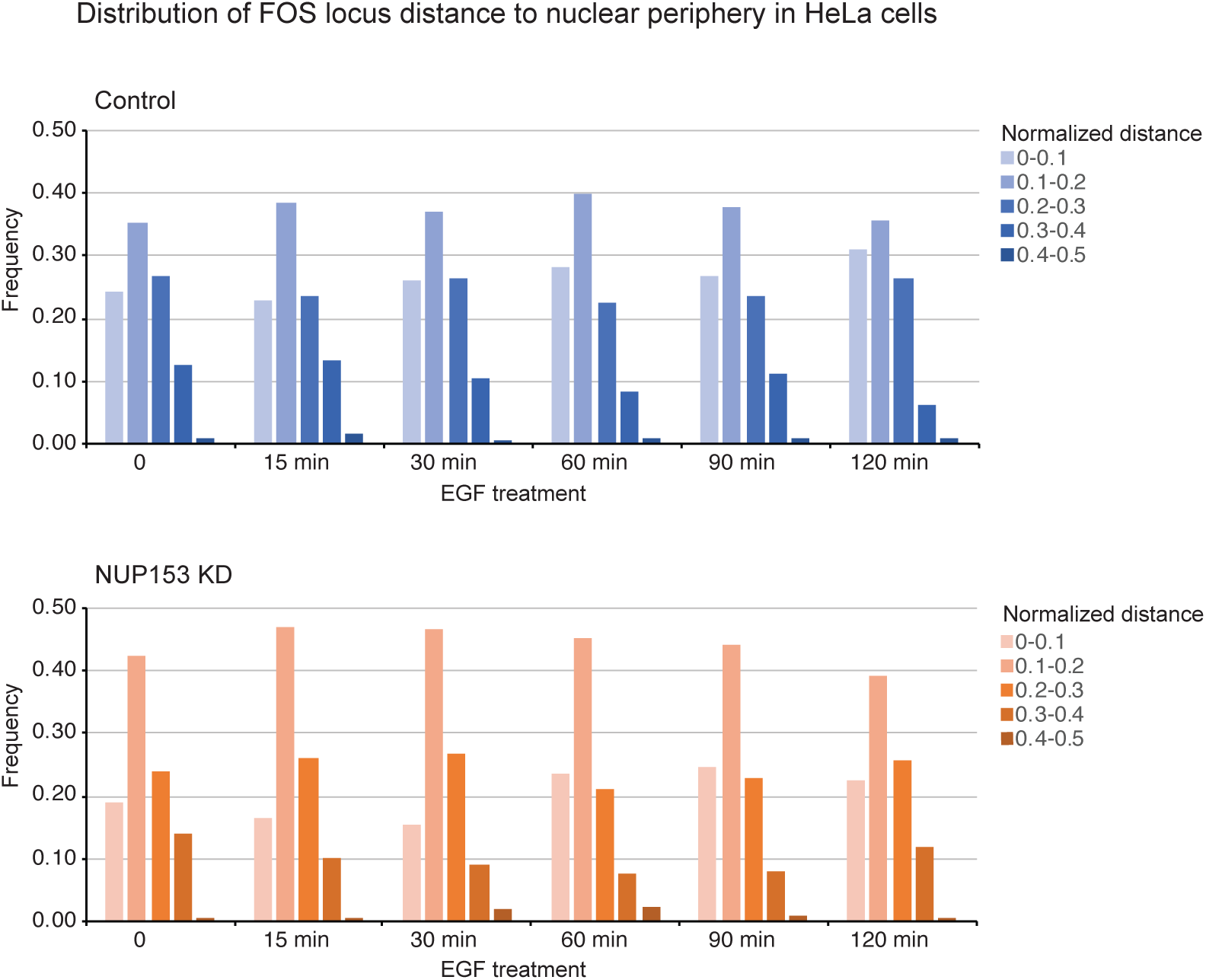
Subnuclear position of *c-FOS* locus with respect to nuclear periphery in HeLa cells. **Related to** Figure 7. Distribution of *c-FOS* locus distance to nuclear periphery in control and NUP153 KD HeLa cells was measured based on DNA FISH at the indicated time points +/− EGF (50ng/ml). Frequencies at a normalized distance (ND) of 0.0-0.5 are shown. ND= *c-FOS* locus to periphery distance/cell diameter (d), where d=(2xnuclear area/π)^0.5^. Control HeLa cells (-EGF, n=182; 15 min EGF, n=186; 30 min EGF, n=150; 60 min EGF, n=146; 90 min EGF, n=181; 120 min EGF, n=139); NUP153 KD HeLa cells (-EGF, n=66; 15 min EGF, n=138; 30 min EGF, n=170; 60 min EGF, n=106; 90 min EGF, n=237; 120 min EGF, n=170). Experiment was repeated twice.

**Table S1:**
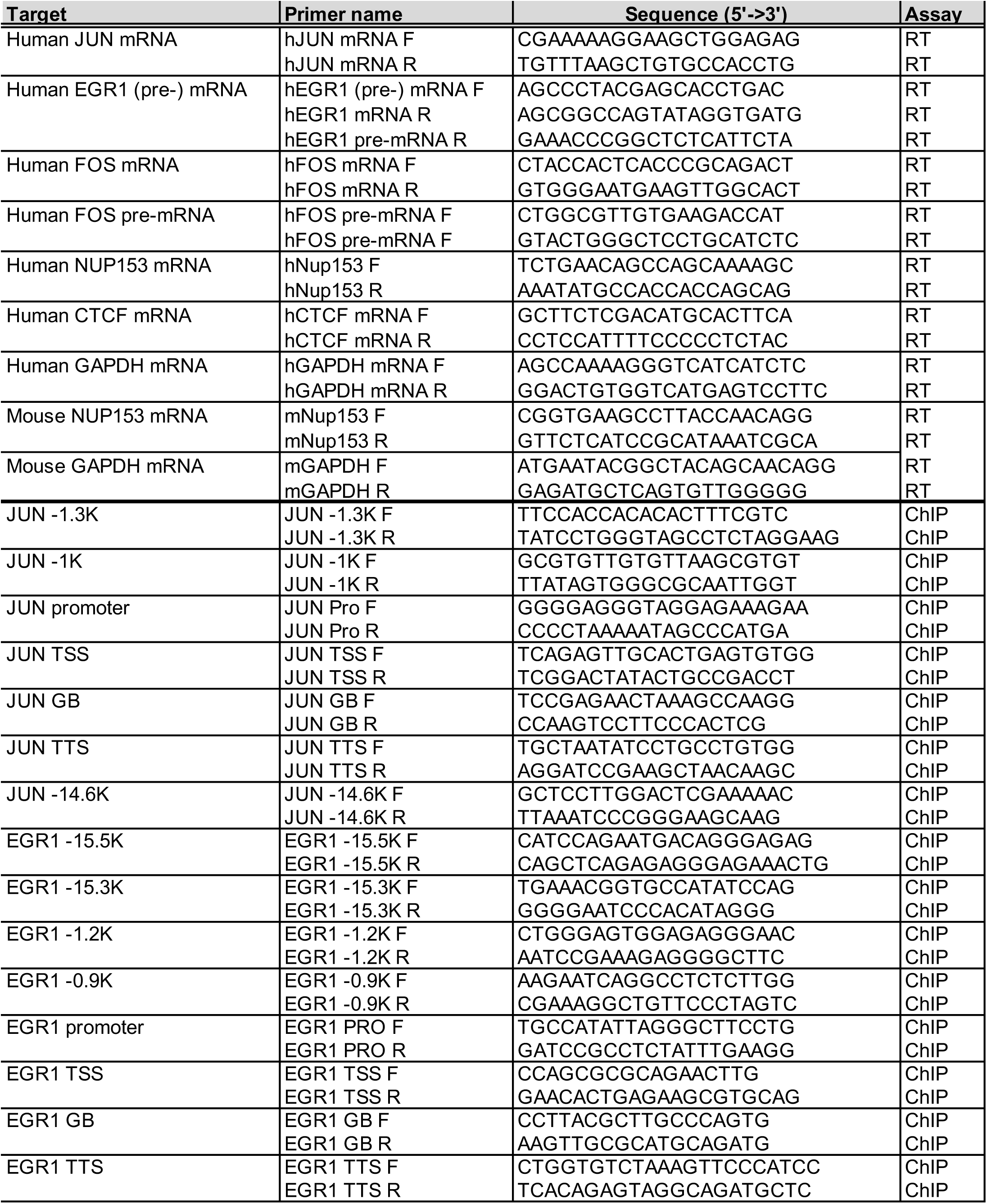
Primer sets used for RT or ChIP real-time PCR analyses, Related to Figure 4, 5, 6, and S6.

